# Learning activator–inhibitor dynamics at the cell cortex with neural likelihood ratio estimation

**DOI:** 10.64898/2026.05.06.722433

**Authors:** Ondrej Maxian, Edwin Munro, Aaron R. Dinner

## Abstract

A key question in cell biology is how cell-scale organization emerges from a given set of molecular players and rules of interaction. Given its multiscale nature, addressing this question requires a combination of experimental perturbation, mathematical modeling, and parameter inference. We leverage recent advances in each of these fields, focusing in particular on neural-network methods for simulation-based inference, to study how cell-scale patterns of Rho GTPase activity are defined by molecular-scale activator-inhibitor interactions with filamentous actin. Using an existing model of this interaction, we demonstrate that an over-expressive, regularized classification neural network can approximate the likelihood of data arising from a particular parameter set. We show that variations in F-actin assembly dynamics can be inferred directly from experimental data, but only if the network is made less sensitive to model misspecification. We use our approach to interpret perturbation experiments in which increasing RhoGAP coexpression increases the frequency and coherence of Rho activity waves in frog eggs. After showing that the known functions of RhoGAP are insufficient to explain experimentally-observed dynamics, we use neural methods to suggest an alternative pathway by which RhoGAP could decrease filament nucleation rates to sustain waves. Our work yields specific, experimentally-testable predictions and illustrates how a combination of traditional forward models and modern inference tools can aid in unraveling mechanisms of self-organization.

## 1 Introduction

In cell biology, a central question is how cell-scale dynamics emerge from microscopic interactions among constituents [1–3]. Given that experimental systems are complex [4, 5], mathematical modeling is often used to correlate differences in molecular parameters with changes in cell-scale patterns [6–10]. As some of the molecular processes driving self-organization can now be directly characterized experimentally [11–15], model parameters can be validated for the first time, making it all the more important to utilize accurate and efficient methods for parameter inference.

Since mathematical modeling involves simplifying choices, there is almost never a one-to-one correspondence between the model and the experimental system at the molecular level. As a result, multiple parameter combinations may fit the data comparably well, and it is useful to quantify how many combinations do so. For this reason, it is more appropriate to infer *distributions* of model parameters, rather than values. Model and experimental outputs are complex, however, and it is usually not possible to write a closed-form mapping from parameters to data. Thus it difficult to assign a probability (or “likelihood”) of observing the data given a particular parameter set (with a probability assumed “prior” to comparison with the data), or a (“posterior”) probability that the data come from a specific set of parameters. The only plausible approach is therefore to use repeated simulations, or “simulation-based” inference (SBI) [16] to compare the model output to the data.

Patterning of Rho GTPases on the cell membrane for cytokinesis, cell migration, and morphogenesis serves as a paradigm of self-organization in development [17–22]. One stunning example is the interaction between active membrane-bound RhoA (hereon Rho) and F-actin/RhoGAP at the cell cortex (Fig. 1A), in which active Rho autocatalyzes its own activation through RhoGEF (Ect2), while simultaneously initiating the assembly of actin filaments (F-actin) through downstream effectors; to complete the activator-inhibitor loop, F-actin recruits RhoGAP (RGA-3/4) to inactivate Rho [23, 24]. Despite conservation of the key molecular players across cell types [23–29], the dynamics of Rho activity in starfish and frog cells (traveling waves) [24, 27] differ sharply from those in *C. elegans* (transient pulses) [23,25,26]. As the dynamics in both cases can be manipulated by perturbing certain components of the circuit (e.g., Rho effectors, RhoGAP) [24, 27], the broader question becomes how each molecular player shapes the overall patterns of Rho activity *in vivo*.

**Figure 1:**
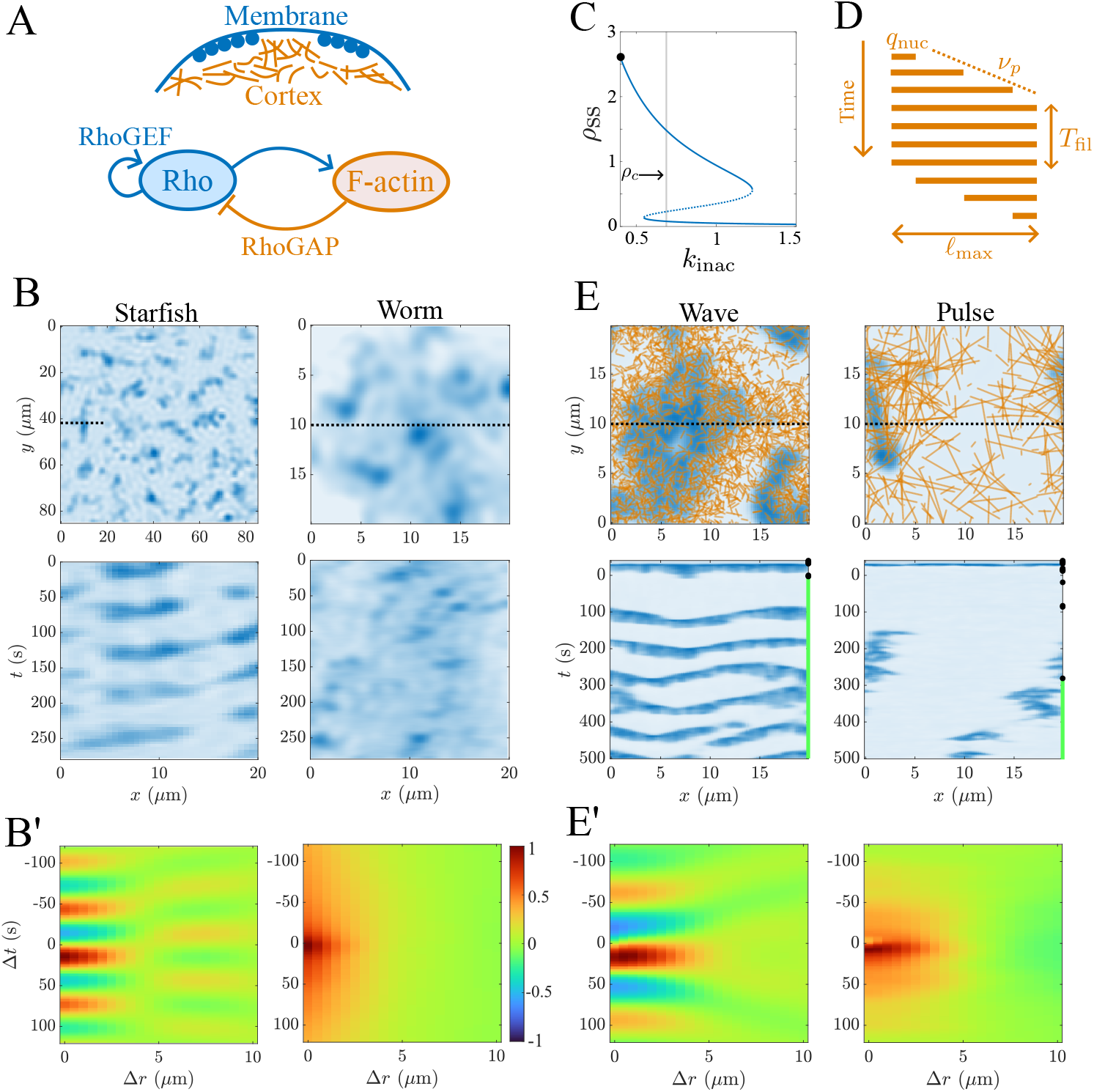
Rho and F-actin form a versatile activator-inhibitor system at the cell cortex, which can be understood using cross-correlation and computational modeling. (A) System schematic. Active membrane-bound Rho signals the assembly of cortical actin filaments, which recruit RhoGAP to inhibit Rho. (B) The dynamics of Rho in starfish [24, Fig. 1C] and worm (*C. elegans*) [26] embryos. Top plots show stills, bottom plots show kymographs along a 20 *µ*m width (the dashed lines). These data are pre-processed to accentuate the Rho signal (see Methods). (B’) Corresponding cross-correlation functions, computed on the filtered data using (1) (colorbar applies to all cross-correlation functions throughout this paper). (C–E) Hybrid model of Rho/F-actin dynamics. (C) Rho is represented as a continuum according to (5a). This diagram shows the steady states in the absence of diffusion with all parameters fixed except 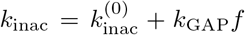. By default, 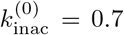 so that the Rho dynamics lie in a bistable regime. (D) F-actin is represented using discrete filaments, which nucleate with rate *q*_nuc_, assemble with speed *ν*_*p*_ to length *ℓ*_max_, remain fixed for time *T*_fil_, and then disassemble with rate *ν*_*d*_ = *ν*_*p*_. These filaments are convolved with a Gaussian to define a continuum field (Fig. S1). (E) Sample snapshots and kymographs of simulations representing waves and pulses. If all excitations extinguish, we reinitiate (force) them by switching from 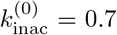 to 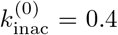 (see panel C and the Methods paragraph on dynamic steady states); switching points are marked by black symbols. (E’) Simulated cross-correlation functions, computed over the longest simulation period without forcing (green lines in kymographs in panel E).

We recently developed a mathematical model which captures the spectrum of behaviors from traveling waves to transient pulses [30]. Our model represents colocalized F-actin and RhoGAP explicitly in filamentous form, imparting the system with a stochasticity and anisotropy that is key to reproducing the behavior observed in *C. elegans*. To infer parameters, we previously employed approximate Bayesian computation (ABC) to approximate the likelihood function by errors in summary statistics [31–33]. In addition to making sampling specific to the data set being studied, this approach prevents full characterization of the likelihood function, instead only identifying the parameter sets that best reproduce the data [16, 34, 35] (see also Section 4.2). Thus, our previous work was not high-throughput enough to examine multiple sets of perturbation experiments, nor accurate enough to compare multiple hypotheses for how pattern variation could emerge.

Modern machine learning algorithms have the potential to revolutionize SBI [16, 36–41], but their rapid development has made it unclear how best to incorporate them in complex inference problems. Neural techniques for inference vary from unsupervised learning of the posterior (e.g., via normalizing flows) [42–45] to supervised learning of likelihood ratios between parameter sets [36]. The past three years alone have seen the application of neural techniques for SBI to synthetic data in economics and finance [46], astrophysics [47, 48], cardiovascular modeling [49], and cell spreading [50]. Extending these approaches to experimental data is more challenging because the data do not directly come from the model (there is some misspecification), which can result in overconfident and incorrect predictions [51–53] (see also Section 4.3). Applications to experimental data have consequently been relatively limited, but do include both static structural data on cell [54–57] and organismal scales [58], as well as dynamic time series data [55, 59–61].

Here we make use of neural SBI to perform high-throughput, accurate inference on our model of Rho GTPase dynamics, making it possible to identify and differentiate hypotheses for how perturbations in molecular players yield cell-scale changes in Rho activity. Our primary innovation is to adopt neural likelihood ratio estimation (NLRE) [34], training a discriminative classifier to learn the mapping from data and parameters to the corresponding likelihood-to-evidence ratio. In the first half of the paper, we validate this approach on both synthetic and previously-studied experimental data [30], in the process demonstrating the sensitivity of posterior estimation to classifier regularization and the dimension of the training data. Our validation also shows how NLRE provides complementary information to errors in summary statistics.

In the second half of the paper, we use our approach to understand how RhoGAP expression levels could tune the frequency and coherence of waves that arise in activated frog embryos [24]. Beginning with a model that accounts for the known interactions in this system [24], we show that increasing GAP activity with other variables fixed does not reproduce the experimental data. Using the robustness of NLRE, we identify two other parameters (filament lifetime and nucleation rate) that could change in response to additional RhoGAP activity. We then use the accuracy of NLRE to infer the more likely of the two, hypothesizing that the amount of F-actin nucleated independently of Rho must decrease as GAP activity increases. This prediction, which does not require any additional simulations once the network is pre-trained, is experimentally testable using fluorescence microscopy and particle tracking [14, 62], and provides an alternative to previous predictions which utilized more traditional approaches [24].

## 2 Self-organization of Rho and F-actin at the cell cortex

The spatiotemporal dynamics of Rho (and F-actin/RhoGAP) can be visualized by imaging fluorescently tagged biosensors which bind active Rho (F-actin/RhoGAP). Using this strategy, transient cortical waves of Rho activity were first observed in starfish and frog oocytes at the beginning of cytokinesis, just after anaphase onset [27]. Quasi-steady versions of these dynamics can be reconstituted in earlier stages of the cell cycle by coexpressing additional copies of RhoGEF (Ect2) and RhoGAP (RGA-3/4) [24, 27], as shown in Fig. 1B.

In *C. elegans*, by contrast, the dynamics of Rho are characterized by transient, meandering pulses of activity [23,25,63]. Previous work has shown that these pulses are *not* a result of contractile instabilities [23, 25], but rather result from the intrinsic dynamics of the activator-inhibitor circuit of RhoA and F-actin. In fact, the pulsatile dynamics in wild-type and myosin-depleted embryos are indistinguishable [26, 30], and so for simplicity we show an embryo depleted of myosin in Fig. 1B (see Methods for pre-processing steps).

### 2.1 Cross-correlation quantifies spatiotemporal activator/inhibitor dynamics

Because the experimental dynamics are stochastic, reproducing individual trajectories of Rho amounts to overfitting noise, and the overall spatiotemporal dynamics are best understood through system-level statistics. We make use of the the cross-correlation between Rho and F-actin, which has been used extensively in previous work [24,27,30] (Fig. 1B’). At time lag Δ*t* and radial distance Δ*r*, the cross-correlation is defined as

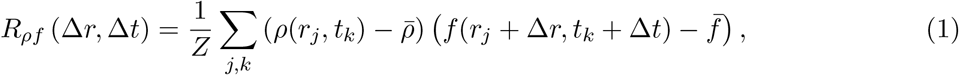

where 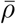 and 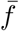 are the mean amount of Rho and F-actin, respectively, and the normalizing factor sets the maximum absolute value |*R*_*ρf*_ | = 1. For the starfish data, the cross-correlation gives information about the temporal frequency and spread of waves, as it is oscillatory with period roughly 60 s (corresponding to the period observed in kymographs), and *R*_*ρf*_ (0, Δ*t <* 0) *<* 0 results when Rho spreads into regions that lack F-actin (making for a negative correlation with F-actin in the past). In *C. elegans*, Rho and F-actin are positively correlated for Δ*t* = 0 and Δ*r* ≤ 3 *µ*m, corresponding to the typical size of a pulse. As we observed previously [30], the cross-correlation in *C. elegans* lacks the signature of traveling waves observed in starfish, and is more characteristic of standing patterns (being roughly constant in time without any negative values).

### 2.2 “Forward model” which reproduces variation across cell types

We previously showed that the differences in Rho activity across cell types can be explained by accounting for changes in F-actin assembly dynamics [30]. Specifically, while in the starfish system F-actin is relatively homogeneous, fluorescent movies of F-actin (LifeAct/Utrophin) in *C. elegans* [23, 26, 64] clearly show a heterogeneous meshwork with visible bundles and holes. Motivated by this, we developed a model which explicitly accounts for the *filamentous* nature of F-actin (Fig. 1C–E) [30]. In this two-dimensional model (Fig. 1(C–E)), active Rho evolves according to a reaction-diffusion equation ((5a) in Methods), where the reaction terms are parameterized to give low and high stable steady states (Fig. 1C). The dynamics of Rho are coupled to a discrete network of actin filaments which nucleate, grow to a fixed length, stay standing for some time, and then shrink (Fig. 1D). To encode the activator-inhibitor interaction, actin filaments are preferentially nucleated in regions of high Rho activity, and F-actin acts as an inhibitor of Rho, with inhibition strength *k*_GAP_ modeling the RhoGAP concentration in the system (see Methods for equations).

To specifically examine how F-actin dynamics dictate Rho activity, we fix the values of all parameters *except* those which determine F-actin assembly and inhibition strength. We reduce F-actin dynamics to one spatial and one temporal parameter by fixing the growth and shrink rates, and scanning over filament length *ℓ*_max_ and lifetime *T*_fil_. Defining parameters *f*_*b*_ and *f*_*ρ*_ as the rates of basal and Rho-mediated filament nucleation (see Methods) gives a total of five unknowns (*k*_GAP_, *ℓ*_max_, *T*_fil_, *f*_*b*_, *f*_*ρ*_) which set the overall model behavior. As a large amount of actin leads to a complete loss of Rho activity, we constrain the amount of F-actin by computing a binary five-dimensional interpolating function *S*(*k*_GAP_, *ℓ*_max_, *T*_fil_, *f*_*b*_, *f*_*ρ*_) which assigns parameter sets which are unable to sustain excitation a “prior probability” (of observing the parameters) of zero (see Methods).

## 3 Neural likelihood ratio estimation (NLRE)

Once the prior and model are defined, the full inference problem comes into focus. Given experimental data ***x***, we would like to predict the probability the data come from the model with parameters ***θ*** = (*k*_GAP_, *ℓ*_max_, *T*_fil_, *f*_*b*_, *f*_*ρ*_). This is the posterior density *p*(***θ***|***x***), which by Bayes theorem is given by

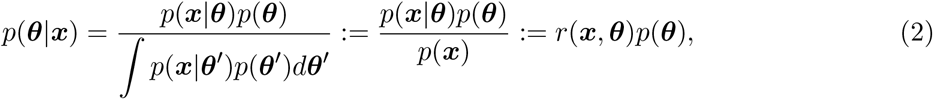

where *p*(***x***|***θ***) is the likelihood of observing the data from a given set of parameters, *p*(***θ***) is the prior probability of observing those parameters, and *p*(***x***) is known as the evidence of the data. Thus, *r*(***x***|***θ***) is the likelihood-to-evidence ratio. We set ***x*** equal to a low-dimensional representation of the cross-correlation function, which is obtained by performing principal component analysis on a set of 10,000 cross-correlation functions (see Methods and Fig. S2). We use the cross-correlation function to represent the data because it is reproducible across experimental and simulated samples (unlike pointwise trajectories), and we compress it to focus the learning on coarse features rather than fine details (which are less robust across replicates).

In approximate Bayesian computation (ABC), the posterior density is estimated by drawing a parameter set ***θ*** ~ *p*(***θ***) from the prior, running the forward model to obtain a simulated data set ***x***_*s*_, and then estimating 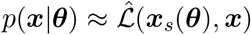 using an approximate likelihood based on summary statistics. In addition to requiring one simulation per sample, this approach does not adapt well to new data, as it requires an additional scan of parameter space for each new data set. By contrast, in NLRE, the likelihood-to-evidence ratio *r*(***x***|***θ***) is approximated by a classification neural network, which is pretrained to pair parameter sets with data. This requires a high upfront cost for training, but a reduced cost thereafter, since the “amortized” likelihood [39] (neural network) is much cheaper to query than the forward model, and can be used for multiple data sets at once without having to resample/retrain.

To perform NLRE, we make use of the so-called “likelihood ratio trick” [34, 36] to approximate *r*(***x***|***θ***) in (2) via a classification neural network. We specifically follow the approach of Hermans et al., who train a classifier on pairs of samples ***x*** and parameters ***θ*** by (somewhat unusually) injecting the parameters ***θ*** as features. The job of the classifier is then to distinguish *dependent* sample-parameter pairs (***x, θ***) ~ *p*(***x, θ***) (label *y* = 1) from *independent* sample-parameter pairs (***x, θ***) ~ *p*(***x***)*p*(***θ***) (label *y* = 0); i.e., separate parameters ***θ*** that generate ***x*** from those that do not (Fig. 2; see Methods for more details on classifier training). Once the classifier is trained, a new data set ***x*** and potential parameter match ***θ*** can be assessed by computing the probability that the data comes from that parameter set (equal to the probability *y* = 1). From this, the likelihood-to-evidence ratio can be approximated as

**Figure 2:**
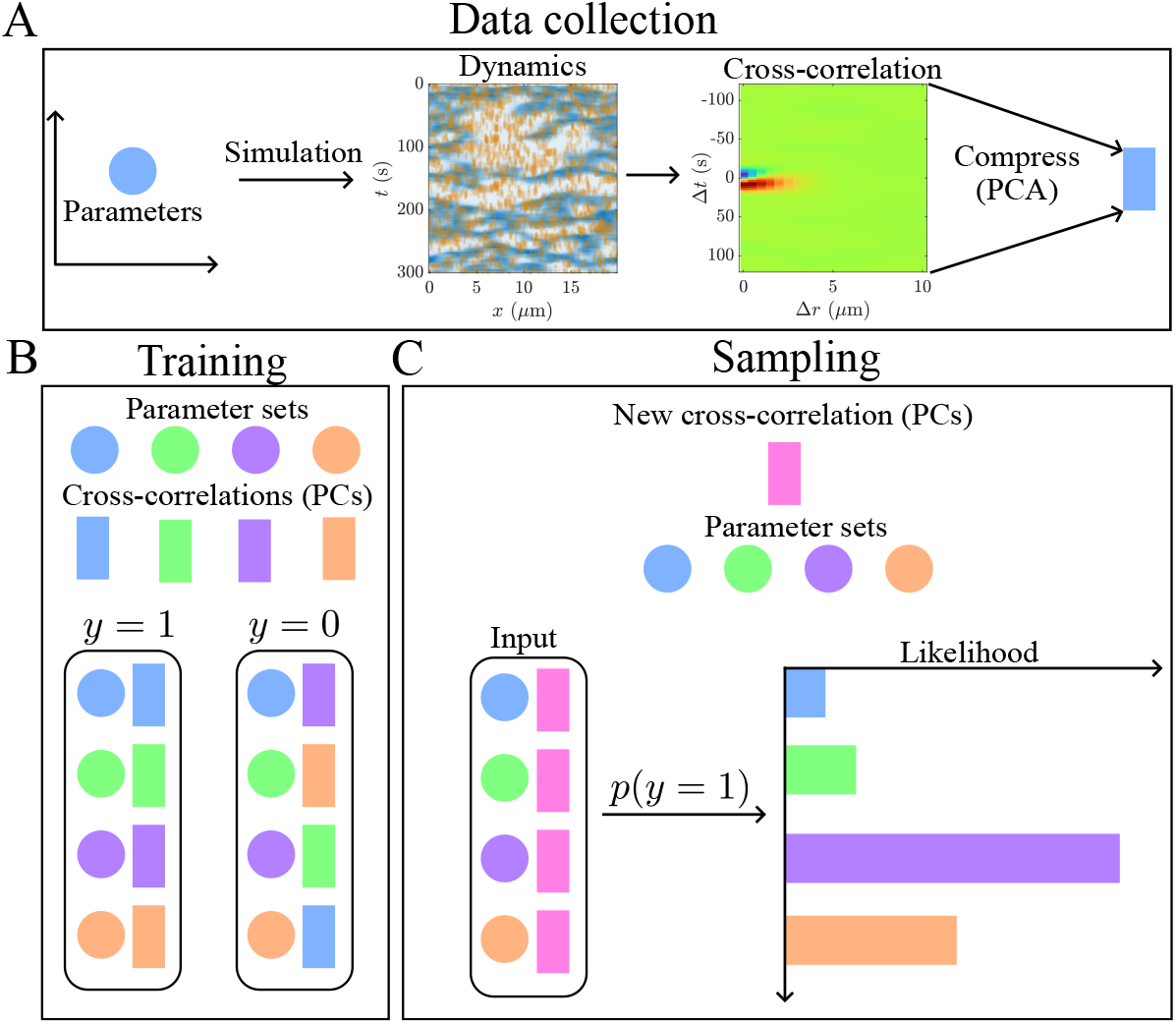
Deploying neural likelihood ratio estimation (NLRE) [34] to estimate the likelihood-to-evidence ratio. (A) To construct a data set, we sample parameter space by running the forward model and computing a compressed version of the cross-correlation for each parameter set. (B) In the training phase, we train the classifier to distinguish between dependent and independent parameter-sample pairs. (C) In the sampling phase, we input a new sample (e.g., experimental data), then compute the probability this sample is associated with each parameter set, from which the likelihood can be computed by (3). In this schematic, the purple parameter set “most resembles” the pink new sample, and so the classifier assigns it the highest likelihood.

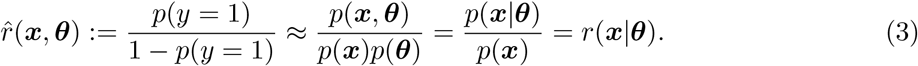

This result is proportional to the posterior density *p*(***θ***|***x***) for parameters with nonzero prior (since the prior is constant where it is nonzero; see Methods for more details on inference). Throughout the paper, we use a hat to refer to quantities *approximated* by the neural estimator.

## 4 Results

Our ultimate goal is to use NLRE to infer the minimal set of parameters responsible for changes in cell-scale Rho activity. Prior to doing this, however, we must be confident that the likelihood ratio estimator 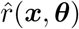 provides a good approximation to the model’s true likelihood function *p*(***x***|***θ***). Thus, we first validate NLRE on synthetic data in Section 4.1 by choosing a parameter set ***θ***^∗^, generating samples from the true likelihood function *p*(***x***|***θ***^∗^), and comparing to the approximate likelihood function 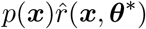 generated by NLRE [34,36]. In Section 4.2, we show further that the likelihood ratio estimate tracks with the error in the cross-correlation function but penalizes errors in some PCA modes more than others. In Section 4.3, we move on to experimental data, finding that additional regularization is essential to prevent overfitting when there is model misspecification. In Section 4.4, we validate the inferred model parameters (with the additional regularization) against kinetic data from *C. elegans* [11, 23, 62]. We also show how NLRE can efficiently survey the posteriors for the starfish and worm data and interpolate between them. Finally, in Section 4.5, we apply our methodology to infer how changing RhoGAP expression alters the frequency and coherence of waves of Rho activity in activated frog oocytes. After demonstrating that existing knowledge of how RhoGAP enters the circuit (Fig. 1A) is insufficient to reproduce the data, we show that there must be negative feedback of RhoGAP activity on basal filament nucleation rates to sustain traveling waves.

### 4.1 Validation on synthetic data reveals importance of classifier architecture and regularization

Following previous work in NLRE [34, 36], we validate the likelihood-to-evidence ratio (LER, (3)) by evaluating the identity 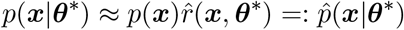 for a fixed parameter set ***θ***^∗^. The distribution *p*(***x***|***θ***^∗^) can be sampled by repeatedly running the forward model with ***θ*** = ***θ***^∗^ to generate sample cross-correlation functions (and representations in PCA space). Meanwhile, samples from *p*(***x***) can be obtained by running the forward model with a new parameter set chosen randomly from the prior at each sample. Weighting these samples by the ratio 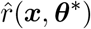 then gives 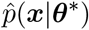. Column 1 of Fig. 3 gives a visual representation of this procedure in the first two components of PCA space: samples from *p*(***x***|***θ***^∗^) are shown using black dots, while samples from *p*(***x***) are colored by the estimated likelihood ratio 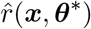. The ratio estimator is accurate when black samples from *p*(***x***|***θ***^∗^) overlap with the highest-probability regions of the likelihood (red points), and when there are no red points outside the support of the black samples. The first part of this section develops some basic metrics to quantify this comparison, while the second part discusses results for specific classifier architectures.

**Figure 3:**
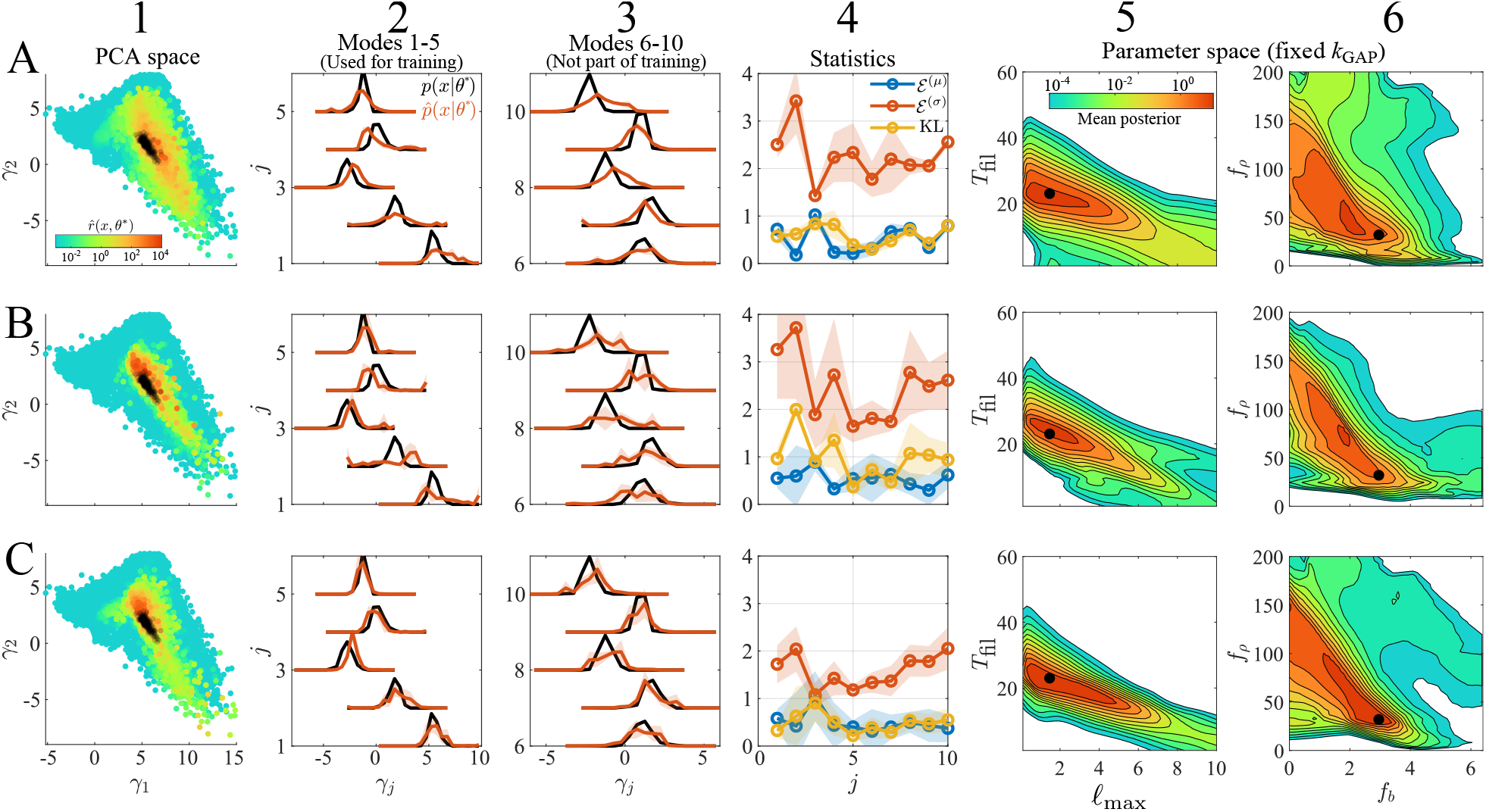
NLRE on synthetic data is most successful with over-expressive, regularized neural networks. Column 1: Samples from *p*(***x***|***θ***^∗^) in the first two components (*γ*_1_, *γ*_2_) of PCA space (black), compared to likelihood ratio estimates 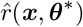 for random samples from *p*(***x***). Column 2: One-dimensional histograms of distributions *p*(***x***|***θ***^∗^) (black) and 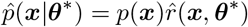 (red) in PCA space for the first five modes used to train the classifier. Column 3: Same histograms, but for modes 6–10, which are not used to train the classifier. Column 4: Summary statistics in (4) comparing the one-dimensional distributions for each mode. The relative error in the mean (blue), ratio of standard deviations (red), and KL divergence (yellow) are shown. Columns 5 and 6: Two-dimensional projections of the posterior *p*(***θ***|⟨***x***(***θ***^∗^)⟩), where the black dot is the true parameters ***θ***^∗^ and *k*_GAP_ = 0.25 is fixed to the true value to make the F-actin densities (*f*_*b*_, *f*_*ρ*_) meaningful. (A) Classifier has one hidden layer of 40 nodes with no *L*^2^ weight penalty. (B) Classifier has three hidden layers of 20, 40, and 20 nodes, with an *L*^2^ weight penalty of *λ* = 10^−5^. (C) Classifier has three hidden layers of 20, 40, and 20 nodes with an *L*^2^ weight penalty of *λ* = 10^−4^.

#### 4.1.1 Metrics for quantifying accuracy of ratio estimator

Previous work compared *p*(***x***|***θ***^∗^) and 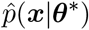 by training a classifier to distinguish between samples from 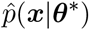 and *p*(***x***|***θ***^∗^) [34, 36]. Paradoxically, success is defined by a *failure* to classify the samples correctly, meaning that the distributions are indistinguishable. In our case, approximating the likelihood is challenging because of randomness in the simulated outputs, which arises from the stochastic nature of the F-actin dynamics (as *in vivo*). Even in local regions of two-dimensional parameter space, significant variation of the first two cross-correlation PCA coordinates is observed (Fig. S3), and the true likelihood function *p*(***x***|***θ***) changes its width depending on the location of ***θ*** in parameter space (for a few examples, see the spread of the black dots in Column 1 of Fig. S4). Thus, we do not think that it is realistically attainable, nor necessary, for the distributions 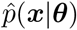 and *p*(***x***|***θ***) to be indistinguishable in our system, and we ask instead for 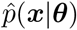 to be a “good enough” approximation to the likelihood function.

To quantify “good enough,” we compute the one-dimensional marginal likelihood over each dimension of PCA space and plot the true marginals together with their approximation in Columns 2 and 3 of Fig. 3. To quantify the comparison in each PCA dimension *γ*_*j*_, we consider the KL divergence of the two distributions, as well as the relative error in the mean (*µ*_*j*_ and 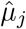, respectively) and ratio of standard deviations (*σ*_*j*_ and 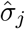, respectively):

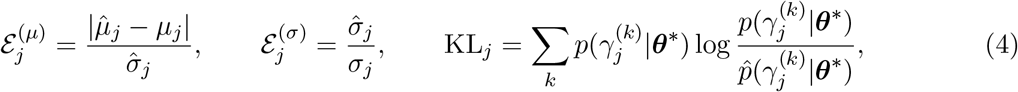

where the KL divergence is computed by summing over discrete 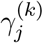 spaced 0.5 apart in PCA coordinate space. To estimate the mean and variance in (4), we construct four different approximations 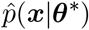 using two different sets of 20,000 samples from *p*(***x***), and two different instantiations of the classifier. The comparison of statistics is shown in Column 4 of Fig. 3.

An alternative way to validate the ratio estimator is to perform inference in the reverse direction. That is, instead of fixing ***θ***^∗^ and checking the probability of different cross-correlation functions ***x***, we could fix 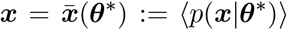 (the mean cross-correlation for ***θ*** = ***θ***^∗^), then compute the likelihood ratio 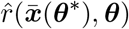 for an arbitrary parameter set ***θ***. For a “good” ratio estimator, the posterior probability 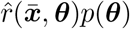 should peak around the original parameters ***θ***^∗^ used to generate 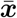. This version of validation is less rigorous because there is no access to a ground truth, as many (potentially unknown) parameter sets could generate cross-correlation functions similar to 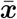 (for instance, halving *k*_GAP_ but doubling the amount of F-actin leaves the dynamics unchanged in the mean field limit). Yet, it is still useful because fixing the data and scanning over the parameters is how the ratio estimator will be used in practice. Therefore, we make the amount of F-actin unique by fixing 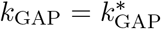 and sample a 50 × 50 × 50 × 50 grid in the four-dimensional space of remaining parameters. We project the approximate posterior 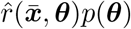 onto two dimensions by averaging over the other parameters and overlay the true parameters ***θ***^∗^ (black dot) with the average posteriors in Columns 5 and 6 of Fig. 3.

#### 4.1.2 Results for different classifier architectures

We now use our metrics to determine the performance of different classifier architectures for our specific model. Because there are only five parameters, we expect the likelihood of a particular parameter set to be determined approximately by the first five PCA modes of the cross-correlation, so we begin by training the classifier on the first five modes only. A fully connected neural network has an input layer with size equal to the number of inputs (size of ***x*** + size of ***θ***, which equals 10 in our case), an output layer with size equal to the number of outputs (the number of classes, which is two here since *y* = 0 or 1), and an arbitrary number of “hidden layers” in between (each of arbitrary size). Our goal in this section is to identify the simplest hidden layer architecture that reduces the error metrics in (4).

We begin with a single hidden layer of 40 nodes. When we train the neural network without an *L*^2^ penalty on the weights, we still obtain overly wide likelihood functions 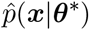, with ℰ^(*σ*)^ ∈ [2, 3] (Figure 3A, Columns 2–4). Previous work [34] showed empirically that these erroneously wide likelihoods can result when the classifier lacks a sufficient number of parameters to fully describe the likelihood function; in other words, the classifier might not be sufficiently expressive. Based on this reasoning, we expand the network to three hidden layers with 20, 40, and 20 nodes. Even when trained with a small *L*^2^ penalty on the weights, this classifier generates estimates for the likelihood which are high variance (note the large error bars in Fig. 3B, Columns 2–4) and have significantly higher KL divergences than the less expressive classifier. An examination of PCA space (Fig. 3B, Column 1) reveals that high variance predictions come from isolated points which are erroneously assigned high likelihood, a classic symptom of overfitting [65–67]. Increasing the *L*^2^ penalty on the classifier weights results in approximations with desirable metrics: KL ≤ 1, ℰ^(*µ*)^ ≤ 1 and ℰ^(*σ*)^ ≤ 2 (Fig. 3, Column 4). Though less rigorous, it is also noteworthy that the true parameter set sits in the region of highest posterior probability only for this last classifier (Fig. 3C, Columns 5–6).

Interestingly, though the classifier is trained only on modes 1–5, the likelihood estimates for modes 6–10 are also good approximations of the truth (Fig. 3C, Column 3), with the errors ℰ^(*µ*)^, ℰ^(*σ*)^, and KL all lower for modes 6–10 than for some of the modes we used for training (Fig. 3C, Column 4). In fact, of the first 40 modes, only mode 28 has a KL divergence larger than 1, and only mode 34 has a standard deviation ratio larger than 3 (not shown). Thus, in the context of synthetic data, getting the first five modes right is sufficient to reproduce the entire spectrum.

Summing up, we find that a regularized, expressive classifier, with only the first five PCA modes used as input, is sufficient to approximate *p*(***x***|***θ***^∗^). Although our approximate likelihood is distinguishable from the true likelihood, the peak in each dimension is at most one standard deviation away from the true peak, and the approximate standard deviation in each dimension is at most twice the true standard deviation. Figure S4 shows that these observations are robust to several different choices of ***θ***^∗^ (different widths of the true likelihood *p*(***x***|***θ***^∗^)), implying that the ratio estimator is robust and accurate for synthetic data.

### 4.2 Likelihood ratio estimate complements cross-correlation error

The main advantage of NLRE is the amortized likelihood function, which allows for inference of parameters from new data without having to perform additional simulations. But how does the approximate likelihood compare to errors in summary statistics? To answer this question, we compare the output of NLRE to errors in the cross-correlation, which we employed in previous work [30].

To compare the two metrics, we sample 5000 cross-correlation functions ***x*** ~ *p*(***x***), compute the mean error relative to 200 samples ***x***^∗^ ~ *p*(***x***|***θ***^∗^) (using the Frobenius norm), and compare this to the mean estimated LER 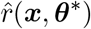 (Fig. 4 shows the same ***θ***^∗^ as Fig. 3; Fig. S5 shows an additional ***θ***^∗^). Generally speaking, parameter sets with low cross-correlation error are deemed high likelihood, while parameter sets with high cross-correlation error are deemed low likelihood (Fig. 4A, S5A). There are, however, a few samples that have high likelihood and high cross-correlation error, or low likelihood and low cross-correlation error. These samples merit further examination, as the inferred likelihood depends on the choice of method.

**Figure 4:**
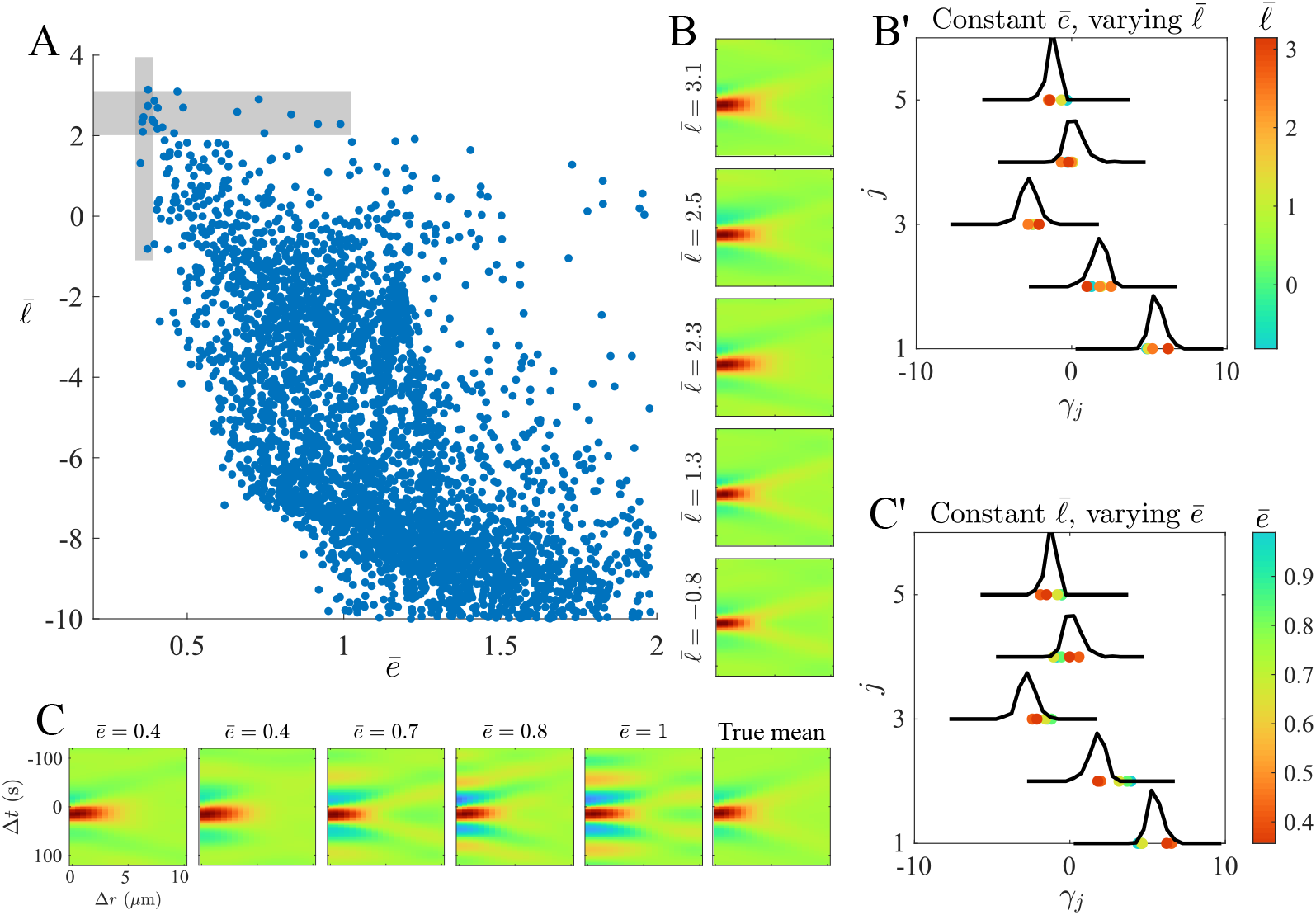
Likelihood ratio and cross-correlation error contain complementary information. (A) Scatter plot comparing the mean error in cross-correlation 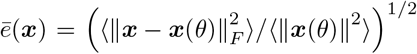 with the mean likelihood ratio 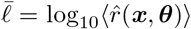. Gray rectangles denote high-probability regions (size 0.1 in cross-correlation and 1 in log likelihood space) where there is variation in one quantity but not the other. (B) Samples of constant cross-correlation error and varying log likelihood. (B’) Corresponding samples in PCA space, colored by the log likelihood, and compared to the true likelihood *p*(*γ*_*j*_|***θ***^∗^). (C) Samples of constant log likelihood but varying cross-correlation error. (C’) Corresponding samples in PCA space, colored by the cross-correlation error, and compared to the true likelihood *p*(*γ*_*j*_|***θ***^∗^). The true mean cross-correlation ⟨*p*(***x***|***θ***^∗^)⟩ is shown at the bottom of (B) and right of (C).

We first isolate samples with low cross-correlation errors and varying estimated likelihood by considering the left-most region (vertical rectangle in Fig. 4A) where cross-correlation error *ē* is roughly constant (at most 10% difference in relative error), but the estimated likelihood 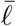 changes significantly (by at least 3 orders of magnitude). Samples of these cross-correlation functions (spaced uniformly in 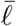) demonstrate subtle changes which move the cross-correlation farther from the true mean as the likelihood decreases (Fig. 4B, S5B). For an oscillatory cross-correlation, the central positive band appears to decrease in width (moving from top to bottom in Fig. 4B), while for a diffuse positive cross-correlation, a negative patch appears near (Δ*r*, Δ*t*) = 0 (bottom panel of Fig. S5B). The change in likelihood is best understood by examining the cross-correlation functions in PCA space (Fig. 4B’), where low-likelihood samples are positioned closer to the edges of the true likelihood distribution (black distributions in Fig. 4B’), with mode 5 appearing particularly sensitive in both parameter sets (Figs. 4B’ and S5B’). Since the fifth mode has the highest peak in the marginal likelihood, we conclude that the classifier penalizes sharply-constrained modes more than others, thus overcoming some of the limitations associated with a single summary statistic (the error).

We next isolate samples with high estimated likelihood and varying error by considering the top-most region (horizontal rectangle in Fig. 4A) where the likelihood estimate 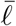 is roughly constant (within one order of magnitude), but cross-correlation error *ē* changes significantly (by at least 0.5). Interestingly, these cross-correlation functions (Fig. 4C and S5C, approximately uniformly-spaced in *ē*), all resemble the original function but typically have more extreme features as the error increases. Indeed, it is possible to increase the error by over 50% while remaining within the support of the likelihood function (*ē* = 0.7 in Fig. 4C’), which suggests that error can be an unnecessarily harsh statistic. On the other hand, very high errors (*ē* = 1 in Fig. 4C’) fall outside of the true likelihood (for mode 2 especially), which implies that the estimated likelihood might be erroneously large in these cases.

Overall then, we can say that cross-correlation errors and estimated likelihoods are *complementary* statistics. Looking only at cross-correlation errors can discount the nonzero width of the true likelihood, especially when some modes are more sharply constrained than others. At the same time, however, abnormally large cross-correlation errors are a sign the likelihood function is erroneously wide.

### 4.3 Inference on experimental data requires additional regularization and higher-dimensional training data

Having validated the model on synthetic data, we now transition to experimental data, first considering the cross-correlation functions from starfish and worms (Fig. 1B’). As we previously used ABC to identify parameter regions matching these data sets [30], this represents a midpoint between synthetic data (where rigorous validation is possible) and completely new experimental data (where we have yet to determine the model parameters). In the context of NLRE, the data (which we denote by capital ***X***_*d*_) are given, and the only quantity that we can evaluate is the likelihood ratio estimate 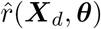 for model parameters ***θ***. Multiplying by the prior then gives the estimated posterior 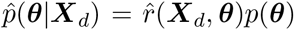. Figure 5(A,E) shows two-dimensional projections of the posterior for (A) starfish and (E) worm data using the optimal classifier settings from the synthetic case.

**Figure 5:**
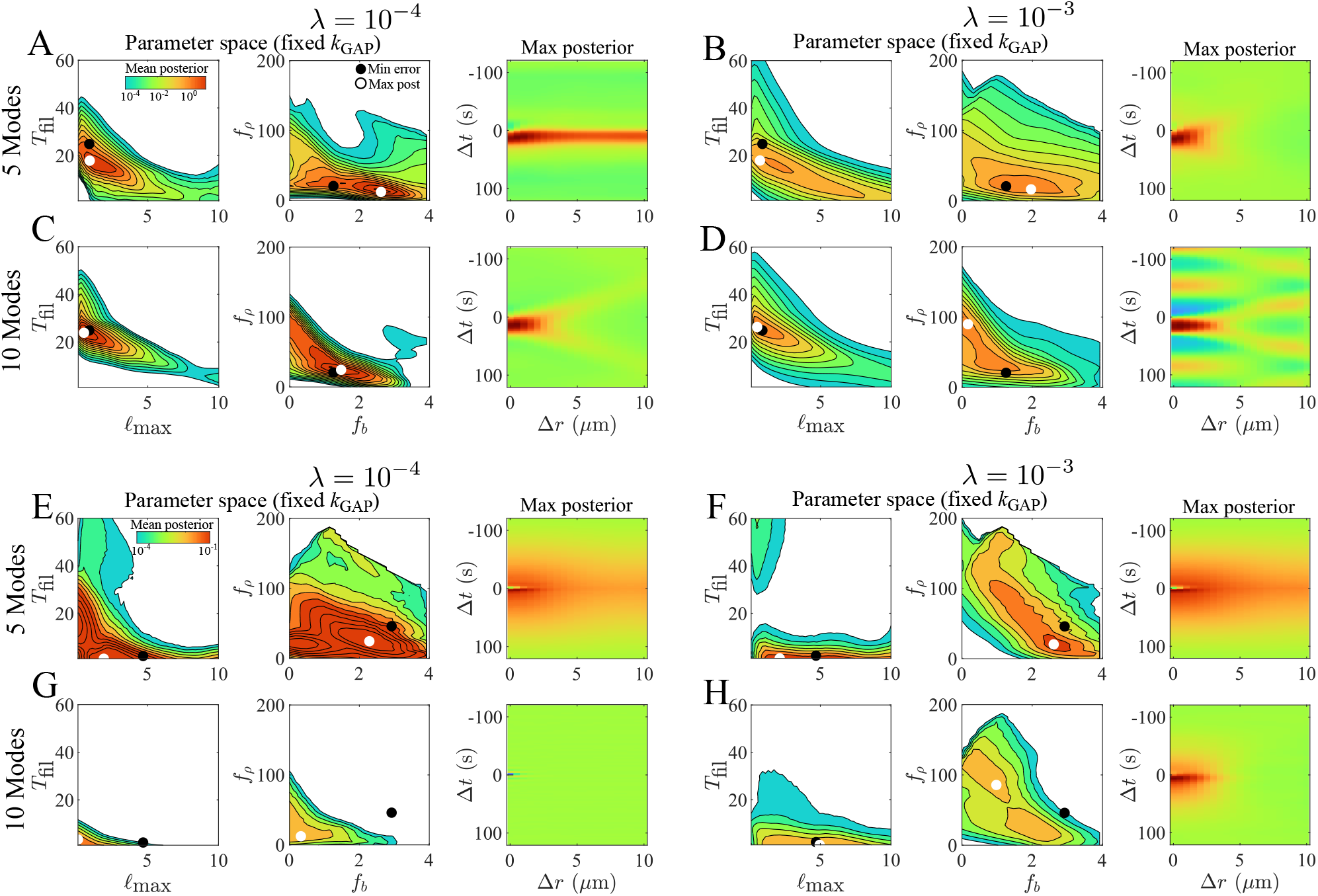
Experimental data require additional regularization strategies due to model misspecification. First two columns of each panel: Two-dimensional projections of the estimated posterior 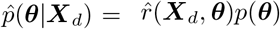 when *k*_GAP_ = 0.4 is fixed and ***X***_*d*_ is the (A–D) starfish or (E–H) worm data (see Fig. 1B). Black dots: parameters that minimize cross-correlation error. White dots: parameters that maximize the posterior (occasionally not at the peaks due to the two-dimensional projection). Third column: The average simulated cross-correlation function obtained using the parameters that maximize the posterior. (A, E) Default classifier parameters (*λ* = 10^−4^, 5 PCA modes) from synthetic validation. (B, F) Additional regularization (*λ* = 10^−3^) with 5 PCA modes. (C, G) More (10) PCA modes, with regularization held at *λ* = 10^−4^. (D, H) More PCA modes and additional regularization (*λ* = 10^−3^, 10 modes).

As the ratio estimator is pretrained on synthetic data, the posterior estimate might be poor when the data ***X***_*d*_ do not resemble the training data. To get a sense of the accuracy of the posterior, we identify (from simulations of 50,000 parameter sets with *k*_GAP_ = 0.4 fixed) the simulated data ***x***_*m*_(***θ***_*m*_) that minimizes ∥***X***_*d*_ − ***x***_*m*_∥, and then plot the corresponding parameters ***θ***_*m*_ on top of the estimated posterior distributions (first two columns of Fig. 5). Since minimal errors should correlate with maximal likelihood estimates (Fig. 4), ***θ***_*m*_ (indicated by the black dot) ought to be near the peak posterior ***θ***_mp_ (indicated by the white dot). As an additional check, we simulate the model twice with ***θ*** = ***θ***_mp_ (two samples of ***x*** ~ *p*(***x***|***θ***_mp_)) to compute an average cross-correlation function (third column of Fig. 5). Ideally, this should resemble the experimental data. In addition to these simpler tests, in Fig. S6 we also define an estimated probability distribution 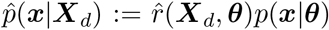 which is based on sampling *p*(***x***|***θ***) for random ***θ***, then weighting the result by 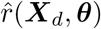. Ideally, 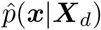 should peak near ***X***_*d*_, or at least near ***x***_*m*_, the model’s closest approximation to ***X***_*d*_.

Unfortunately, directly applying the same classifier architecture and settings used for synthetic data to experimental data yields a posterior which fails to satisfy these tests (Fig. 5A,E). For both the starfish and worm data, the minimum-error parameter set falls at least an order of magnitude away from the peak posterior, and the peak posterior parameters generate cross-correlation functions that do not resemble the corresponding experimental ones. This behavior contrasts with our analysis of synthetic data and thus must result because the physical model, which enters through the training data, does not capture the true data-generating process (*in vivo* dynamics). This is defined as *model misspecification* in the SBI literature [51, 53, 68]. Since the minimum-error cross-correlation functions ***x***_*m*_(***θ***_*m*_) lie close to experimental data in PCA space (Fig. S6), the model can be made to resemble the starfish and worm data in certain parameter regimes [30]. Thus, the issue is not with the model but with the neural estimator’s ability to find those parameter regimes given experimental data.

We consider two non-exclusive possibilities for why the estimator performs poorly under model misspecification: failure of extrapolation and lack of parameter identifiability. A failure of extrapolation can result when the classifier estimates 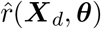 for data outside the training set even if there is no overfitting inside the training set. This phenomenon manifests as high variance in the estimated likelihood ratios (see the shading and blue arrows in Fig. S6A,E), and can be mitigated by increasing the regularization parameter *λ* in the classifier loss function. On the other hand, a lack of identifiability occurs when the modes used are insufficient for the classifier to relate experimental data to training data. This can occur because the PCA modes are computed using *simulated* data and thus need not be representative of experimental data. In this case, parameters can be made more identifiable by increasing the number of modes used for inference.

Figure 5 shows that the combination of more regularization and more PCA modes in training yields reliable posterior estimates for both the starfish (A–D) and worm (E–H) data. In both parameter and PCA space, more regularization alone gives higher relative probabilities of the minimum-error parameter sets, but does little to exclude cross-correlation functions that do not resemble the data (Fig. 5B,F; see also Fig. S6B,F which shows overly-wide distributions 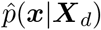 in PCA space). Alternatively, increasing the number of modes alone sharpens the posteriors and makes the parameters identifiable but still gives erroneously-high posterior estimates in some regions of parameter space (Fig. 5C,G; see also Fig. S6C,G). Combining more regularization with more PCA modes gives the best outcomes: posteriors assign high probability to the minimum-error parameter sets, and maximal-posterior parameters generate cross-correlation functions that resemble the data. In PCA space (Fig. S6D,H), the probability distributions 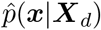 peak near the data and/or its best model approximation (compare for example green arrows in Fig. S6B,D). Interestingly, the minimum-error parameter set for the worm data falls far from the maximum posterior (black dot in Fig. 5H), but the minimum-error cross-correlation function is also far from the data in the first PCA mode (see the blue star in Fig. S6H), so we do not view this as a shortcoming of the ratio estimator. Summing up, the optimal classifier training settings (*λ* = 10^−3^ and 10 PCA modes) combine more regularization (to address overfitting when ***X***_*d*_ is not part of the training data) with more modes (to sharpen identifiability). Figure S7 shows that using these classifier settings on synthetic data widens the estimated marginal likelihoods in some cases, without inducing strong changes in the location of the peaks or KL divergence relative to the true likelihoods.

### 4.4 Validated NLRE reveals how changing F-actin assembly parameters shifts dynamics from waves to pulses

We previously showed that the difference between the worm and starfish data can be explained by choosing parameters in diametrically opposite regions of F-actin parameter space, with starfish having long-lived, short F-actin, and worms having short-lived, long F-actin [30]. A more complete understanding of the transition between the two types of behavior can be obtained by using NLRE to sample the complete parameter space, in this case by fixing *k*_GAP_ = 0.4 and sampling the posterior on a 50 × 50 × 50 × 50 grid in space of *ℓ*_max_, *T*_fil_, *f*_*b*_, and *f*_*ρ*_ (Fig. 6A shows the average posterior in two-dimensional subsets of parameter space). The posterior estimates clearly show that the two parameters of F-actin *architecture* (*ℓ*_max_ and *T*_fil_) are most distinguishable across data sets. Interestingly, the likelihood ratios for the worm data are significantly smaller than for the starfish, which reflects the relative impacts of model misspecification discussed previously (Figs. 5 and S6). Despite this, independent kinetic measurements of F-actin dynamics in *C. elegans* have found maximum filament lengths of 6 *µ*m [62] and subunit turnover times *T*_sub_ = *ℓ*_max_*/ν*_*p*_ + *T*_fil_ on the order 8–10 s [11] (which translates to *T*_fil_ = 2 s). These direct parameter measurements fall squarely within the high-likelihood regime of our inference (blue dot in Fig. 6A), thus validating our ability to predict the microscopic dynamics responsible for pattern formation.

**Figure 6:**
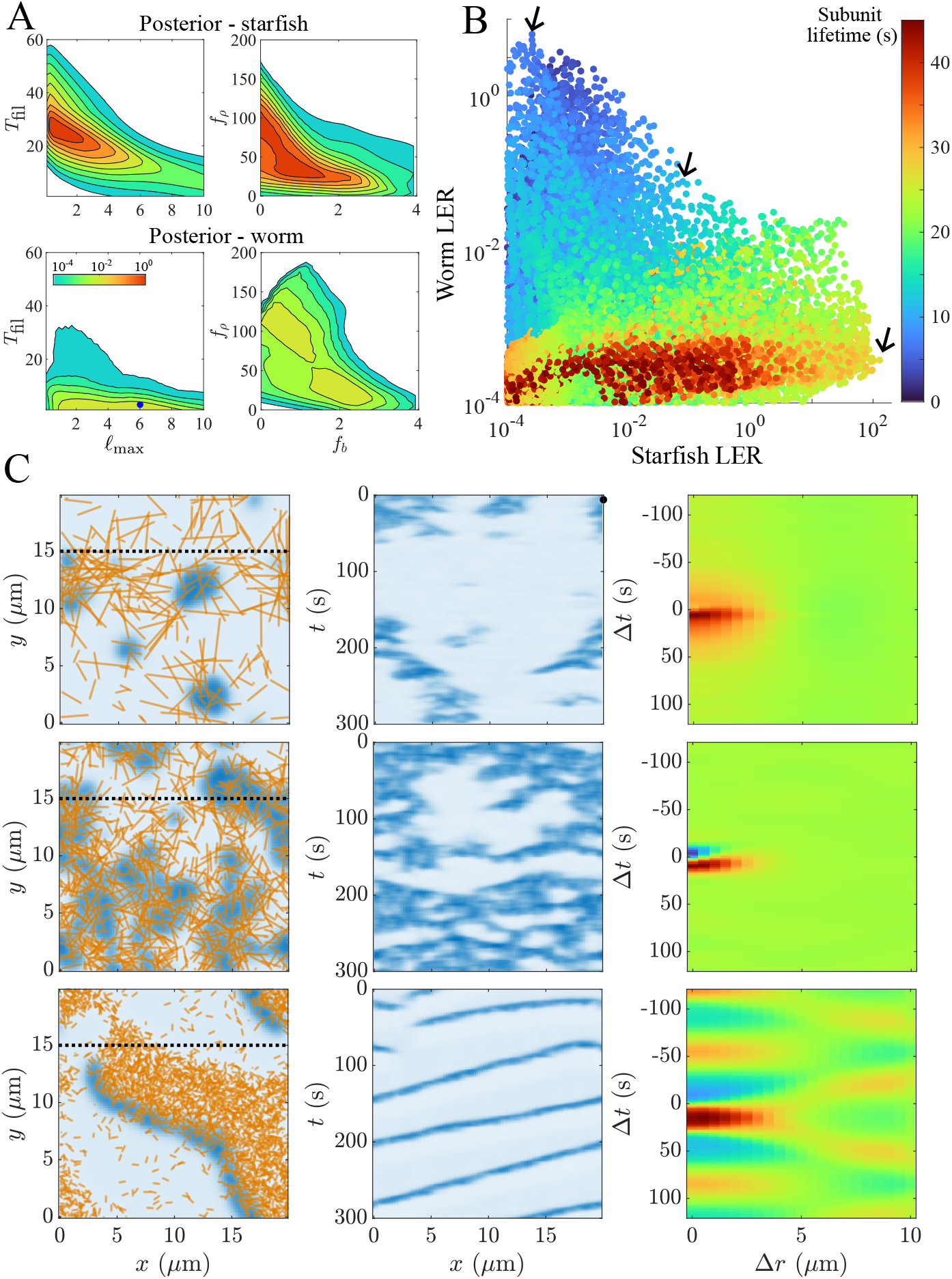
Posterior distributions reveal continuous transition between starfish and worm data, depending on the actin subunit lifetime. (A) Two-dimensional projections of the posterior for starfish (top) and worm (bottom) data with fixed *k*_GAP_ = 0.4 (sampled at 50^4^ parameters). (B) Scatter plot of the likelihood-to-evidence ratio (LER) 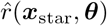 against 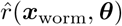 for 10% of the samples (number reduced for clarity, samples taken only at points away from the boundary of the prior, measured as probability ≥ 75% of sustaining excitation). (C) Sampled dynamics at the arrows indicated in (B), from top to bottom. Left panels show snapshots at *t* = 60, middle panels show kymographs about *y* = 15, and right panels show cross-correlation (c.f. data in Fig. 1B’).

Reducing the parameter space to subunit lifetime alone further accentuates the distinction between starfish and worms, and allows us to find parameter sets that interpolate between the two. Plotting the likelihood-to-evidence ratio for starfish 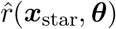 against that for worms 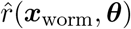 and coloring points by subunit lifetime (Fig. 6B) shows that subunit lifetimes larger than 20 s are favorable for the starfish data alone, while subunit lifetimes smaller than 10 s are favorable for the worm data alone. Simulations using the maximum likelihood parameter set for each data set strongly resemble the original data (Fig. 6C), allowing us to assert with renewed confidence that changes in F-actin assembly dynamics drive the change from Rho waves in starfish to Rho pulses in *C. elegans* [30]. Wave-like dynamics correlate with short, long-lived filaments, which lag spreading Rho wavefronts. Pulse-like dynamics correlate with long, short-lived filaments, which polymerize rapidly out of excitations, allowing them to spread only slightly before being extinguished.

A key advantage of NLRE is that we can classify any parameter set as more wave- or pulse-like, depending on the comparison of likelihood against the two types of experimental data. Figure 6B shows the dividing line between wave and pulse-like data occurs around subunit lifetime of 15 s. To visualize the dividing line, we find a value of *c* = 10^−1.12^ where there is exactly one parameter set with both 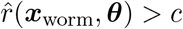 and 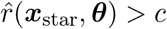 (middle black arrow in Fig. 6B). A corresponding simulation from this parameter set (middle panels of Fig. 6C) shows an interpolation between the wave and pulse-like patterns. The dynamics still have the signature of wave-spread, with negative cross-correlation on Δ*t <* 0, but there are no long-time oscillations in the cross-correlation, which is more similar to the worm data. We emphasize again that the parameter sets in Fig. 6 are *not* part of the training data. Thus, NLRE is able to predict new dynamics which resemble worms, starfish, and in between. This allows us to be more confident in parameter inference when there appears to be a mixture of waves and pulses, as in the data we consider next.

### 4.5 Inferring how RhoGAP could modulate wave dynamics

An important advantage of NLRE is the amortized likelihood function; since the classifier is already pre-trained, we can infer parameters of new experimental dynamics without having to perform more simulations. NLRE therefore provides an opportunity to examine data from perturbation experiments which do not lend themselves to straightforward interpretation. In a typical perturbation experiment, the expression of one of the circuit’s molecular players is modified (e.g., through RNA interference or microinjected RNA), and the resulting dynamics are read out on the cell scale. Such a perturbation can be challenging to interpret because a molecule can play multiple roles, and feedbacks make them hard to parse. Our approach provides an opportunity to infer how the observed patterns could result from changes to specific model parameters, which we can then connect to the molecular players under scrutiny.

We use as an example data collected from activated frog ooctyes [24], where extra Ect-2 (RhoGEF) and RGA-3/4 (RhoGAP) are coexpressed from microinjected mRNA: Ect-2 at constant level (200 ng/*µ*L) and RGA-3/4 at varying levels. Representative trajectories for RGA-3/4 levels ranging from 33 to 1000 ng/*µ*L (Figure 7A) reveal a continuous increase in the frequency and coherence of waves as RhoGAP expression levels increase (data for 0 ng/*µ*L are shown in Fig. S8) [24]. Previous continuum-level modeling reproduced many of the features of these dynamics by incorporating molecular noise (modeled as a spatially and temporally correlated Gaussian stochastic field) into the F-actin disassembly rate [24]. Our approach is to ground this molecular noise in our model of F-actin assembly dynamics, then use NLRE to infer how changes to model parameters lead to the observed changes in wave propagation.

**Figure 7:**
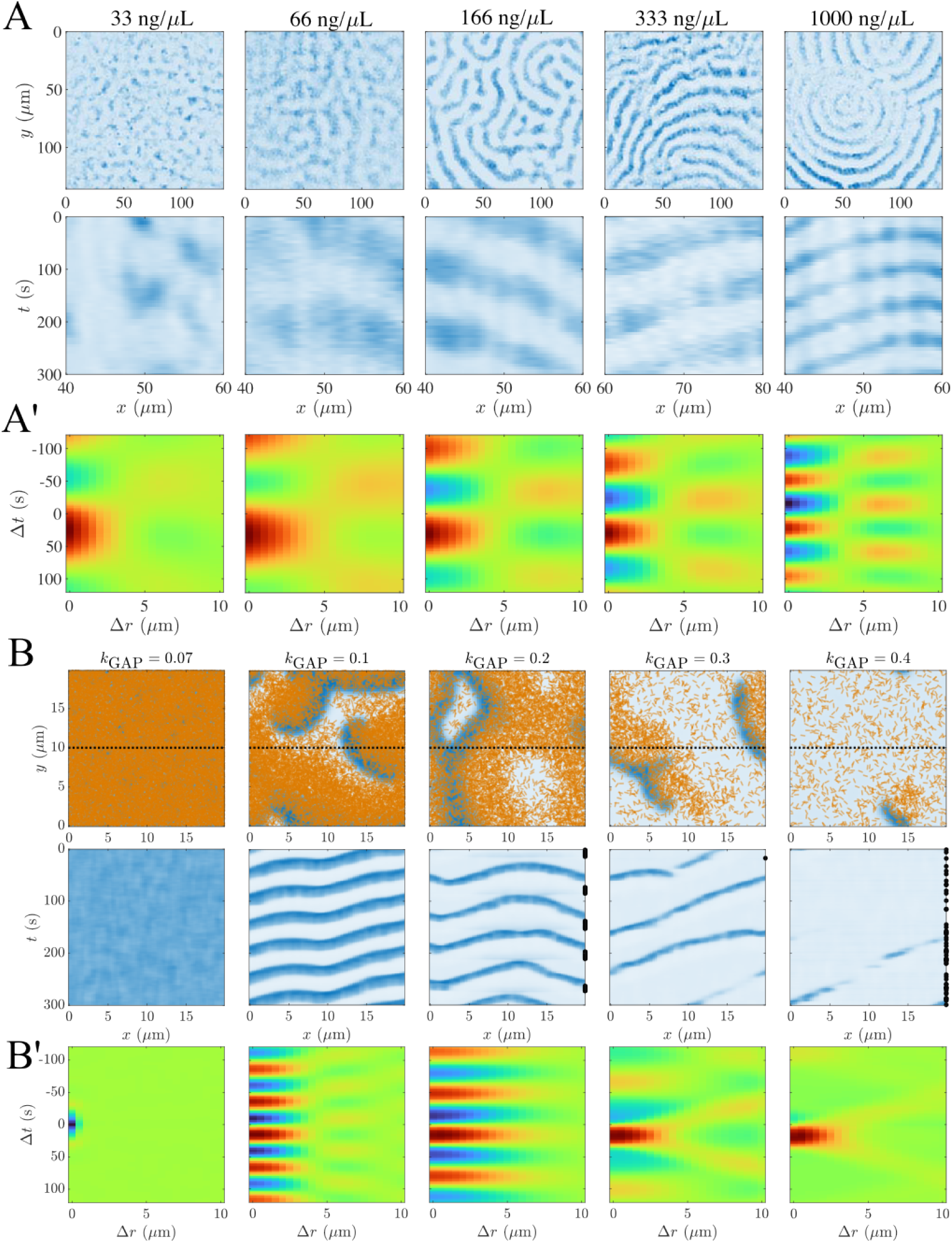
Increasing RhoGAP coexpression leads to faster, more coherent waves *in vivo*, which cannot be explained by increasing GAP strength alone in the model. (A) Snapshots and kymographs (along 20 *µ*m distance) in activated frog eggs coexpressing 200 ng/*µ*L Ect2 and varying levels of RhoGAP (RGA) (indicated in the titles). (A’) Corresponding cross-correlation functions. (B) Model simulations with increasing (from left to right) GAP strength *k*_GAP_ and constant filament assembly dynamics and nucleation rates (*T*_fil_ = 30 s, *ℓ*_max_ = 0.5 *µ*m, *f*_*b*_ = 1 *µ*m^−1^, *f*_*ρ*_ = 80 *µ*m^−1^). (B’) Corresponding cross-correlation functions. In the model, highly coherent waves emerge suddenly once the GAP strength surpasses a threshold, after which coherence decreases as the GAP strength increases.

#### 4.5.1 Increasing GAP strength alone cannot explain the data

Given that only the GAP expression levels are varied in the data, a sensible first step is to fix the parameters of F-actin assembly and increase the GAP strength *k*_GAP_ in the model. When the GAP strength is too low, the F-actin/RhoGAP levels are not high enough to push Rho activity to the lower steady state, and Rho activity levels are high everywhere (left panel of Fig. 7B). A slight increase in *k*_GAP_ destabilizes the uniform state, giving rise to coherent waves with sharp, oscillatory cross-correlation (second panel from left, Fig. 7B). As the GAP strength increases further, the waves become less coherent as stronger inhibition pushes Rho down to its lower steady state in more places. When the GAP strength is sufficiently high, Rho activity ceases unless actively restarted (right panel of Fig. 7B).

As the F-actin length is small (0.5 *µ*m in Fig. 7B), it is not surprising that our results match those of previous continuum models [24]. Similar to those results, the dynamics of Rho below a critical point are dominated by fluctuations around a uniform state. This is supported by the data without any RhoGAP coexpression, though the cross-correlation in that case comes from averaging the random fluctuations, and is consequently more difficult to interpret (Fig. S8). Above the critical *k*_GAP_, waves emerge, and further increases in *k*_GAP_ beyond this point only lower the maximum wave amplitude and temporal width (defined as the time between half-maxima for a given point in space); c.f. [24, Fig. 7B–D].

At the same time, however, analysis of cross-correlation functions (in cases when they are informative, Fig. 7A’ and B’) shows that the model does not fully reproduce the data. In the data, waves emerge gradually, with *more* coherent waves as GAP strength increases, while in the model the waves emerge suddenly, then become *less* coherent as GAP strength increases. Furthermore, in the data, the wave frequency (as seen in the cross-correlation) increases as GAP expression increases, while in the model the wave frequency is approximately constant as GAP strength increases (the central band in the cross-correlation has approximately the same width, regardless of GAP strength). This pattern holds not just for the illustrative (fixed) F-actin assembly parameters in Fig. 7B, but also for F-actin assembly parameters chosen to maximize the likelihood while ensuring that the mean inferred *k*_GAP_ increases as RGA expression levels increase (Fig. S9). Thus, the assumption that RhoGAP levels affect only the strength of inhibition and not any of the other model parameters cannot be correct. This conclusion is not model-dependent: if filament assembly rates are the same, adding additional RhoGAP should only extinguish Rho activity, not sharpen it.

#### 4.5.2 NLRE predicts an essential role for nucleation rates

Given that changing only the GAP strength cannot reproduce the data, a logical next step is to look for another parameter that must change as the GAP strength increases. This also provides an opportunity to leverage the power of NLRE; since the likelihood is a function of all five parameters, we can easily fix a subset and examine which parameters must change to maintain a high likelihood.

There are four candidate parameters that could change alongside GAP strength: the filament lifetime *T*_fil_, the filament length *ℓ*_max_, and the basal and Rho-mediated nucleation rates *q*_*b*_ and *q*_*ρ*_ (which we report in terms of nucleated amounts *f*_*b*_ and *f*_*ρ*_). For each candidate parameter, we identify potential fixed values of the other parameters (*ℓ*_max_, *q*_*b*_, and *q*_*ρ*_) by imposing two constraints: first, with those values fixed, each data set must have at least one (*k*_GAP_, *T*_fil_, *ℓ*_max_, *q*_*b*_, *q*_*ρ*_) combination where the data favor the parameters (likelihood-to-evidence ratio larger than 1). Second, the mean *k*_GAP_ (measured via the first moment of the posterior distribution) must be an increasing function of RhoGAP expression levels (see Methods and (7) for more details). We plot the mean posterior in two-dimensions, averaged over all potential values of the three fixed parameters, then choose a specific parameter set that balances high posterior probability with increasing mean *k*_GAP_ values (via maximizing (8)). For this parameter set, we plot the corresponding two-dimensional posterior over the varying parameters (Fig. S11), then simulate with parameters chosen from the highest likelihood regions (and such that *k*_GAP_ and the other varied parameter vary monotonically with RhoGAP expression).

This procedure identifies two candidate parameters that could be changing with *k*_GAP_, with all other candidates failing to satisfy our two criteria. A first possibility is the filament lifetime *T*_fil_, which could decrease with increasing RhoGAP levels (Fig. 8A). While it is possible to see waves of increasing coherence in this case (Fig. 8C), the optimal *k*_GAP_ varies over a small range, from *k*_GAP_ = 0.3 for the lowest coexpression levels (33 ng/*µ*L) to *k*_GAP_ = 0.35 for the highest ones (1000 ng/*µ*L). Thus, an increase of 30-fold RNA corresponds to only a 17% increase in *k*_GAP_. Simulations with representative parameter sets (Fig. 8C) suggest that, when inferred *k*_GAP_ levels do not vary much, NLRE is focusing on common features of all five cross-correlation functions, and the increases in the mean *k*_GAP_ with RGA coexpression are coincidental. Based on these considerations, as well as a Bayes’ factor calculation which shows a relatively low probability for this model (Methods and Fig. S10), we disfavor a mechanism in which *T*_fil_ alone changes with *k*_GAP_.

**Figure 8:**
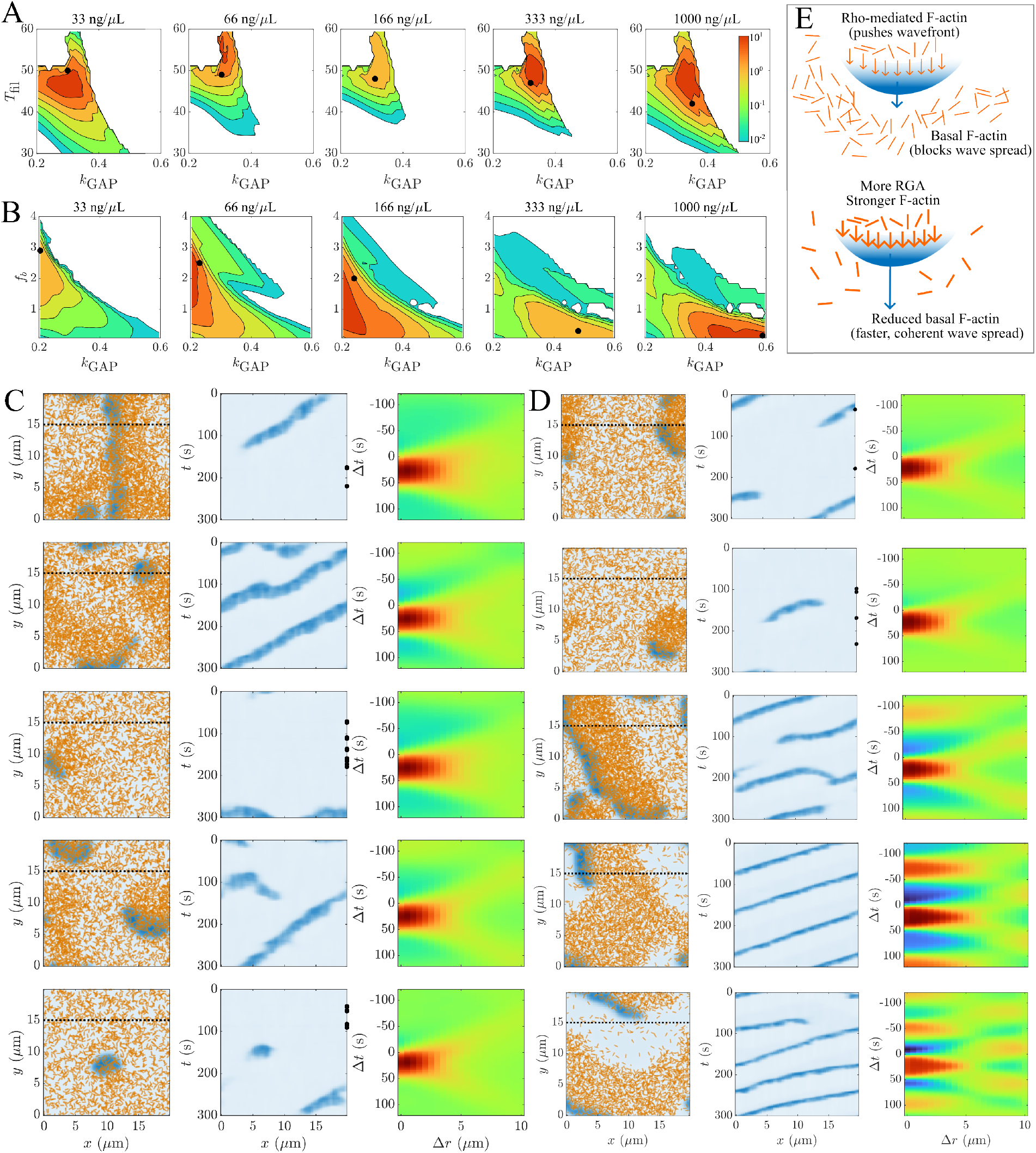
To reproduce data, basal F-actin assembly must be reduced as GAP strength increases. (A, B) Average posteriors for each coexpression level when exactly one parameter is varied in addition to *k*_GAP_. (A) Varying *T*_fil_ alone in response to *k*_GAP_ gives a narrow band of plausible parameter values. Black dots show samples in (C), taken for a specific set of fixed parameters which maximize the posterior while also respecting increases in *k*_GAP_ (see (8) and Fig. S11). (B) Varying the basal assembly rate *f*_*b*_ gives a rich distribution of probability which shifts appropriately as expression levels increase. Black dots show samples in (D). Sample dynamics for the varying lifetime model (GAP strength increases from top to bottom; fixed parameters: *ℓ*_max_ = 0.50, *f*_*b*_ = 0.096*ℓ*_max_(2*ℓ*_max_ + *T*_fil_), *f*_*b*_ = 1.1*ℓ*_max_(2*ℓ*_max_ + *T*_fil_)). (D) Sample dynamics for the varying basal nucleation model (GAP strength increases from top to bottom; fixed parameters: *T*_fil_ = 44, *ℓ*_max_ = 0.50, *f*_*ρ*_ = 78). (E) Mechanistic diagram explaining the role of Rho-mediated F-actin (assembles behind Rho wavefront and sets direction of propagation) and basal F-actin (assembles in the bulk, blocking wave spread). Waves spread faster and become more coherent if basal levels decrease when RhoGAP levels increase.

The second parameter that satisfies our criteria is the basal nucleation rate *q*_*b*_ (and by extension the amount of basal filaments *f*_*b*_). In this case, the inferred parameter sets span a broad range in (*k*_GAP_, *f*_*b*_) space, with the smallest expression levels clustering around (*k*_GAP_ = 0.2, *f*_*b*_ = 2), and the largest around (*k*_GAP_ = 0.6, *f*_*b*_ = 0). The remaining fixed parameters (*T*_fil_ = 44 s, *ℓ*_max_ = 0.5 *µ*m, and *f*_*ρ*_ = 78/*µ*m is a representative example) are consistent with our existing knowledge of the frog system; FRAP experiments show F-actin recovery rates on the order 40 s [69], and filaments must be short and homogeneous to generate waves [30]. Simulations along the most likely path in (*k*_GAP_, *f*_*b*_) space reproduce many aspects of the data: waves become more coherent, higher frequency, and have a lower temporal width, and cross-correlation functions become more oscillatory in time, with the central positive band shrinking in size as RhoGAP coexpression levels increase (though not as significantly as in the data). Thus, the model clearly suggests that sharper waves of Rho activity occur when basal nucleation rates are also reduced in response to increasing RhoGAP activity.

Mechanistically, the output of our inference can be understood in terms of the role of basal and Rho mediated F-actin (Fig. 8E). Typically, Rho-mediated actin assembles behind wavefronts of Rho (and stays there for quite some time), effectively “pushing” the wavefront in one direction. Basal F-actin, by contrast, inhibits spreading excitation, blocking the formation of coherent traveling waves. In the model, a high amount of basal F-actin is necessary to reproduce the incoherent waves that arise at low RhoGAP levels. As RhoGAP levels increase, inhibition becomes more intense, and so holding the basal assembly levels constant makes it impossible for waves to maintain their structure, much less become more coherent. Basal filament levels must therefore decrease with RhoGAP. In the case when Rho-mediated nucleation levels are constant, increasing RhoGAP makes the trailing inhibition more powerful, which pushes the wave at a faster speed, decreasing the temporal width in a given location.

#### 4.5.3 Potential molecular mechanisms to tune nucleation

The result that basal nucleation must decrease with RhoGAP expression suggests a number of possible molecular mechanisms, depending on how we interpret basal filament nucleation. In the simplest interpretation, all of the F-actin is still Rho-mediated, and basal assembly occurs from local fluctuations in Rho density, which are not included in the model (and are averaged over in the cross-correlation analysis). RhoGAP would then act to suppress these fluctuations, without altering the rate of Rho-mediated nucleation. Alternatively, basal filaments could be interpreted as F-actin nucleated independently of Rho via a basal rate of formin activity. In this case, RhoGAP would inhibit formin directly (as has been shown in other systems [70]) and Rho-mediated nucleation would also likely decrease with increases in RhoGAP. Finally, interpreting basal nucleation as independent of the Rho pathway yields the hypothesis that RhoGAP inhibits other GTPases and their effectors (e.g., CDC-42/WASP/Arp2/3 [71, 72]). If the amount of F-actin on the cell cortex is conserved [73–75], a reduction in CDC-42-mediated F-actin could cause an increase in Rho-mediated F-actin. Given that the dynamics in Fig. 8D can also be reproduced when Rho-mediated assembly decreases or increases in response to RhoGAP (Fig. S12), these hypotheses would be best differentiated by performing live-imaging of formin molecules, which would yield local rates of Rho-mediated and formin-mediated nucleation.

## 5 Conclusion

In this work, we applied novel methods for parameter inference to learn how the activator-inhibitor interaction between Rho and F-actin dictates cell-scale pattern formation. Our methodological contribution was to use neural likelihood ratio estimation (NLRE) [34] to quantify the correspondence between models and data via classification neural networks. Using the high-throughput likelihood function, we mapped how varying Rho activity patterns could emerge from changes in F-actin assembly dynamics in starfish, worm, and frog cells. We found that the change from pulse-like behavior to cohesive waves could be accomplished by making actin filaments shorter and longer lived [30], and we validated the inferred parameters from NLRE by comparing to *in vivo* kinetic measurements from *C. elegans* [11, 62]. We also found that more cohesive waves could be achieved with higher RhoGAP levels, but only when basal rates of F-actin assembly are reduced, and suggested experiments to test this prediction.

On the methods side, our work revealed the advantages of neural SBI, while also demonstrating the challenges that arise when trying to apply these methods to noisy biological data. Even for low-dimensional representations of the data, our synthetic model exhibited marginal likelihoods with relatively large variance in some regions of parameter space and small variance in others. We showed that an over-expressive classifier, with loss function regularized to prevent overfitting, can approximately learn this space, with approximate marginal likelihoods that are at most twice as wide the true ones. The approximate likelihood provides complementary information to the errors in the cross-correlation function, as it penalizes some aspects of the error more than others. By leveraging the amortized likelihood function, we used 10^5^ samples of the model to predict the posterior for 50^5^ = 3.1 × 10^8^ samples, which gives each sample an effective power of over 1000. Still, 10^5^ samples is quite large, and it will be important to implement some of the recent “cost-aware” improvements to neural SBI as our models move beyond just Rho and F-actin [38, 44, 76–78].

When we attempted to apply NLRE directly to experimental data, we found that even more regularization was required to avoid overconfident posteriors which often arise from model misspecification [52,79]. Recent work has addressed this issue by developing a model for the misspecification, potentially based on a small set of measurements where the true parameters are known, then inverting the error model prior to inferring the likelihood [51, 53]. This approach could be promising here given that kinetic data are increasingly available in organisms like *C. elegans* [11,62]. Of course, developing a better model of the dynamics which reduces misspecification (perhaps by incorporating more molecular-scale features) is a direction to pursue simultaneously.

While we always visualized the model dynamics in concert with the cross-correlation (e.g., Fig. 8), only a compressed version of the latter was actually used to perform inference, and we found that the number of included modes influenced the results. This raises the question: what information is lost when reducing complex dynamics to summary statistics? For instance, while experimental measurements are subject to noise, there is also randomness in the true dynamics that is averaged over when computing the cross-correlation function. A helpful next step would therefore be to optimize the representation of the data, for instance using feature extraction with neural networks [60]. Though not strictly necessary, it would be ideal if the compressed representation of data were still interpretable. A straightforward way to ensure this is to augment the cross-correlation function with additional summary statistics designed to detect other kinds of spatiotemporal phenomena, similar to how the cross-correlation function is optimal for detecting traveling waves,

On the biology side, our main contribution was to show how increasing RhoGAP levels could give rise to waves of increasing coherence and frequency in activated frog eggs. Similar to previous work [24], we first showed that increasing *k*_GAP_ (the model’s GAP strength parameter) can transition the dynamics from a high uniform steady state of Rho (with small fluctuations) to waves with oscillatory cross-correlation functions (Figs. 7 and S9). Unlike in the data, in the model increases in *k*_GAP_ only suppress this peak excitation, as stronger filaments make waves less coherent and ultimately nonexistent (Figs. 7 and S9). Previous work [24] interpreted the disagreement with the experimental data as evidence for saturation of *k*_GAP_ with increasing amounts of microinjected RNA, which could occur if expression levels are limited by translation, or by crowding along actin filaments (see Fig. 8G in [24], where the GAP strength parameter is limited to small values for frog data). In this interpretation, the full spectrum of dynamics is explained by a noisy transition from a uniform steady state to a bistable state with propagating waves and saturated *k*_GAP_. By contrast, our NLRE analysis found that *k*_GAP_ need not saturate, so long as increases in *k*_GAP_ are paired with *systematic* decreases in the basal nucleation rate. In this interpretation, coherent waves are sustained by reducing the amount of inhibition (F-actin) *in the bulk*, which allows waves to propagate without disruption (Fig. 8E). We offered several hypotheses for how these dynamics could arise molecularly, including the possibilities that RhoGAP suppresses fluctuations in Rho density, or that it directly inhibits formins or other GTPases. In this way, our combination of modeling and neural inference yielded specific, testable predictions that were overlooked previously.

Since our inference procedure is now high-throughput, there is no limit to the amount of experimental data we can analyze—including knockdown experiments that examine how accessory proteins such as anillin and septin [80, 81], or anterior PARs [82, 83] impact GTPase activity by modulating F-actin assembly dynamics. Inference will only be useful if the model includes the data, and so learning the full scope of dynamics requires accounting for myosin-mediated contractility, which is also downstream of Rho [26, 84, 85]. When paired with neural SBI, our model will determine how a balance of reaction, diffusion, and contractility shapes patterns of Rho GTPases, and how each of these is in turn modulated by the key cytoskeletal molecular players.

On a broader level, our use of neural SBI to “learn” the relevant mathematical model and parameters provides a roadmap for future work across other self-organizing systems. Whether in cells [86, 87], tissues [88, 89], or ecosystems [90], connecting small and large scales requires a combination of versatile mathematical models and tools for parameter inference. Neural techniques can identify both relevant models and parameters faster than ever before, but it remains unclear how best to deploy them with experimental data. Our work is a step toward answering this question.

## Data availability

Code for simulations, data analysis, and inference is available at https://github.com/omaxian/RhoActinRD

## Acknowledgments

We thank Gilles Louppe for providing details on how to validate the likelihood ratio, Bill Bement and Ani Michaud for providing the data on activated frog oocytes, and Alex Mogilner and Alan Lindsay for feedback on the manuscript. OM thanks Daniele Schiavazzi for suggesting neural methods for SBI. Simulations were carried out on the Research Computing Center Midway cluster at the University of Chicago, and the Center for Research Computing cluster at the University of Notre Dame. OM acknowledges partial support from the Physics Frontier Center for Living Systems (CLS) Spark Fellowship, funded by the National Science Foundation (PHY-2317138); ARD and EM acknowledge support from the CLS as well as the “From Cells to Organs with AI” Initiative at the University of Chicago.

## Methods

### Data analysis

Similar to previous work [30], we perform pre-processing to better visualize experimental data and extract cross-correlation. In all movies, we first extract a square domain in the middle of the oocyte/embryo. For starfish and frog data, we perform difference subtraction, subtracting the last frame from the others, to remove static Rho signal [24, 27]. For starfish and *C. elegans*, we also correct for photobleaching by setting the mean intensity of each frame to be equal to the global mean. For *C. elegans* (where the signal-to-noise ratio is significantly lower), we smooth the signal in time by applying a moving average filter with a width of 20 frames (12 s). For all movies, we remove high-frequency noise in space by applying a Fourier filter to retain the 20 (for *C. elegans*) 50 (starfish), or 100 (for frogs) lowest-frequency modes. In all cases, cross-correlation functions are computed on the filtered data (although the cross-correlation functions on the raw data are similar).

### Mathematical model

#### Governing equations

The dynamics of Rho in our model are governed by the two-dimensional PDE

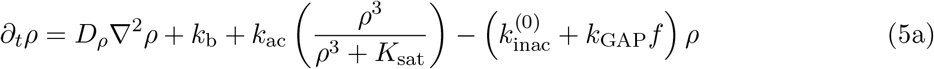

with periodic boundary conditions. Equation (5a) accounts for basal Rho activation and auto-catalytic excitation (through the cubic Hill function), as well as basal Rho inactivation and enhanced inactivation by an inhibitor *f*, which represents colocalized F-actin/RhoGAP [23, 24]. As justified in [30], our model only considers the active form of Rho, as experimental evidence [81, 91] has suggested that inactive Rho is not significantly depleted.

At each point in space, the concentration of Rho sets a nucleation rate of individual actin filaments,

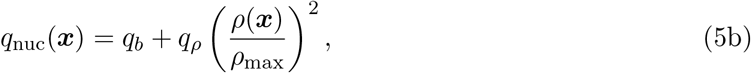

where *ρ*_max_ is the maximum steady state of Rho in (5a) when *f* = 0. Upon nucleation, each filament is assigned a random direction of growth on the unit circle, along which it polymerizes with rate *ν*_*p*_ up to length *ℓ*_max_ (see kymograph in Fig. 1D). Once the filament reaches the maximum length, it remains fixed in space for time *T*_fil_, after which it depolymerizes from the older end with rate *ν*_dp_. The continuum field *f* (***x***) is then defined by convolving the discrete filament centerline positions with a Gaussian point spread function (Fig. S1).

As the nucleation rates are not easily measurable experimentally, we convert these quantities into approximate densities of basal (*f*_*b*_) and Rho-mediated (*f*_*ρ*_) F-actin per domain area by setting *f*_*b*_ = *q*_*b*_*ℓ*_max_*/k*_diss_ and *f*_*ρ*_ = *q*_*ρ*_*ℓ*_max_*/k*_diss_, where the effective filament turnover rate is defined as

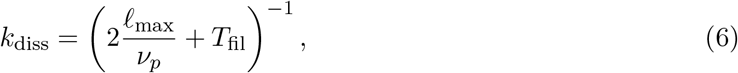

and the factor of 2 accounts for growth and shrinkage. For basal nucleation, *f*_*b*_ is a good measure of the filament density produced independently of Rho, while for Rho-mediated nucleation, *f*_*ρ*_ expresses the filament density that would result if *ρ* = *ρ*_max_ uniformly.

#### Parameterization

We previously showed [30] that the dynamics of Rho must be in a bistable regime where Rho excitations can be initiated via diffusive flux from neighboring regions (e.g., traveling waves), but not automatically *de novo* in the absence of F-actin. We use these observations to constrain the parameters 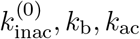, and *K*_sat_ in (5a). Fixing *k*_b_, *k*_ac_, and *K*_sat_, we choose 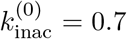, so that in the absence of F-actin (*f* = 0), the total inhibition 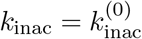 is such that there are two stable steady states of Rho (Fig. 1C). The diffusivity of Rho (*D*_*ρ*_ = 0.1 *µ*m^2^/s) is fixed in accordance with experimental data [23].

In previous work [30], we demonstrated that the dynamics of F-actin are effectively governed by two parameters: one having to do with the spatial extent of growth and the other the timescale of filament turnover. We therefore reduce the dynamics to one spatial and one temporal parameter by fixing *ν*_*p*_ = *ν*_dp_ = 1 *µ*m/s (which corresponds to polymerization speeds in a variety of cell types [62, 92]), and scanning over *k*_GAP_, *ℓ*_max_, *T*_fil_, *f*_*b*_, and *f*_*ρ*_. Table S1 summarizes the fixed and varied parameters.

#### Dynamic steady state and statistics

When there are two steady states of Rho in the absence of F-actin, a uniformly high F-actin concentration could extinguish *all* of the excitation, causing Rho to sit at the lower steady state everywhere. This prevents a quasi-steady state, since an absence of excitation at one time implies an absence of excitation for all time. To induce a dynamic steady state, our approach is to define a threshold Rho concentration, *ρ*_*c*_, equal to the arithmetic mean of the saddle point and higher steady state when 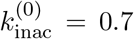 (arrow in Fig. 1C). When the concentration is everywhere below this level, we instantaneously bump 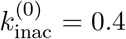 everywhere (black circle in Fig. 1C), until at least one point of Rho activity exceeds the threshold level (in kymographs of Rho activity, e.g., Fig. 1E, black dots denote regions of time when 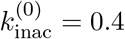). Starting from an initial condition where Rho is at the lower steady state everywhere, we track statistics over the longest self-sustaining simulated interval (the green regions in Fig. 1E).

#### Prior on varied parameters

To approximate the prior *p*(***θ***), we train a discriminative classifier to identify the region in five-dimensional (*k*_GAP_, *ℓ*_max_, *T*_fil_, *f*_*b*_, *f*_*ρ*_) parameter space which is able to sustain excitation (by sampling uniformly within the ranges in Table S1 and recording whether two out of two samples of the model are able to sustain excitations for 120 s or longer). “Sampling from the prior” then amounts to sampling uniformly from the five-dimensional space (*k*_GAP_, *ℓ*_max_, *T*_fil_, *f*_*b*_, *f*_*ρ*_), then eliminating any parameters with a prior value of zero. The prior for the remaining parameter sets is uniform. As the boundary of being able to sustain excitation is complex and not fully resolved by running only 2 trials per parameter set, our prior will sometimes be inaccurate (classifier has 95% validation accuracy). In general, the marginal posterior distributions do not depend on the threshold used to assign a nonzero prior. However, in cases where we are interested in visualizing high-likelihood simulations (e.g., Fig. 6), we reduce false positives by only considering parameters where the score is 0.75 or higher (75% chance of being able to sustain excitation).

### Compressed cross-correlation functions

As discussed in Section 2.1, pointwise trajectories of Rho and F-actin are of little use, since reproducing them often amounts to reproducing noise. One workaround to this is to use a black-box-style approach, where the pointwise trajectories ***x*** are passed to an autoencoder which is trained simultaneously with the likelihood estimator, so that statistics are learned naturally [39, 93–95]. In our case, this presents some challenges because the size of a single movie is enormous (on the order of 100 MB), and because it makes the output less interpretable. Our approach is instead to leverage the cross-correlation function (Section 2.1), which we already know is a good representation of the full spatiotemporal dynamics. However, the two-dimensional matrix of cross-correlation functions (Fig. 1B’) can be arbitrarily large, depending on the spatial and temporal resolution of the experimental data, and so we still require a means of reducing the problem dimensionality.

To compress the cross-correlation while still retaining as much information as possible, we interpolate each function at Δ*t* = −120, −118, … 120 s and Δ*r* = 0, 0.5, 1, … 10 *µ*m (so that the cross-correlation is a 121×21 matrix), then use the classical method of principal component analysis (PCA) to convert each cross-correlation function to PCA space (Fig. S2). To compute the principal components of cross-correlation, we first build a matrix of many cross-correlation functions (where each one is stacked as a single column vector)

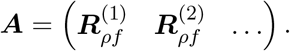

The SVD ***A*** = ***U* Σ*V***^*T*^ gives a list of “components” (the columns of ***U***) and scores **Σ*V***^*T*^. Compressing the cross-correlation into a subset of modes then amounts to setting 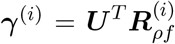 (to obtain the scores of a single cross-correlation), then extracting the first *c* components of ***γ*** (denote this *c* × 1 vector as ***γ***_*c*_). The image can then be decompressed by multiplying 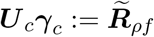, where ***U*** _*c*_ is the matrix containing the first *c* columns of ***U***.

### Training the classification neural network

To train the neural likelihood ratio estimator in our context, we perform the following set of steps:

1. Sample uniformly over parameter space to obtain a set of parameters ***θ***_*i*_ and simulated data sets ***x***_*i*_ (Fig. 2A).
2. Remove any parameter sets from the list where at least one of the runs does not sustain excitation. The number of remaining parameter data pairs is denoted by *N*_*p*_.
3. For parameter set ***θ***_*i*_, input to the classifier (***x***_*i*_, ***θ***_*i*_, *y* = 1) and (***x***_*i*_, ***θ***_*j*_, *y* = 0) for *j* ≠ *i* (Fig. 2B). Repeat this for all *N*_*p*_ parameter sets, so that there are a total of 2*N*_*p*_ observations passed to the classifier.
4. Train the classifier. We use the `ClassificationNeuralNetwork` object in Matlab R2024b with standardized predictors and an iteration limit of 15,000. As noted in the main text, we vary the size of the hidden layer and the regularization parameter in the cross-entropy loss function, which is minimized by the BFGS algorithm.

In step 1, we choose a total of 2 × 10^5^ parameter sets, where for each parameter set we obtain two cross-correlation functions. Thus the total initial number of parameter-data pairs is 4 × 10^5^. Removing parameter sets where excitation is not sustained for both simulations gives *N*_*p*_ = 1.4×10^5^ pairs.

### Using the classification neural network for inference

Given new data ***x***_*n*_ we perform the following set of steps to estimate the posterior *p*(***θ***|***x***_*n*_) (assuming all five parameters could vary):

1. Sample 50^5^ parameter sets by uniformly gridding five-dimensional parameter space over the ranges shown in Table S1.
2. Use the trained classifier to predict whether these parameters will sustain excitation. Assign prior *p*(***θ***) = 0 and posterior *p*(***θ***|***x***_*n*_) = 0 for any parameters predicted not to sustain excitation.
3. For each parameter set ***θ***_*i*_ with *p*(***θ***_*i*_)≠ 0, input (***x***_*n*_, ***θ***_*i*_) to the classifier.
4. The output of the classifier is the probability *y* = 1, which gives 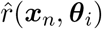 by (3) (Fig. 2C). Assign the posterior 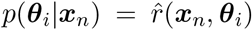 for these parameter sets (the constant prior is normalized out in the next step).
5. To compute the marginal posterior in two dimensions (e.g., over the first two parameters), set

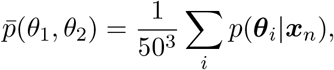

where the sum is over all ***θ***_*i*_ with first two components equal to *θ*_1_ and *θ*_2_.

### Two-dimensional posteriors for frog data

To identify a potential variable *θ*_*v*_ that changes with *k*_GAP_, with the other three variables ***θ***_*f*_ fixed, we scan over ***θ***_*f*_ by sampling a 50 × 50 × 50 grid in the regions of parameter space given in Table S1. For each ***θ***_*f*_, we compile a two-dimensional posterior *π*(*k*_GAP_, *θ*_*v*_, ***θ***_*f*_) by scanning over 50 × 50 values of (*k*_GAP_, *θ*_*v*_). We say that the fixed parameters ***θ***_*f*_ are allowable if

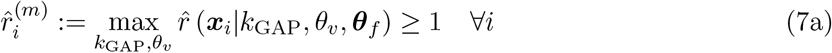

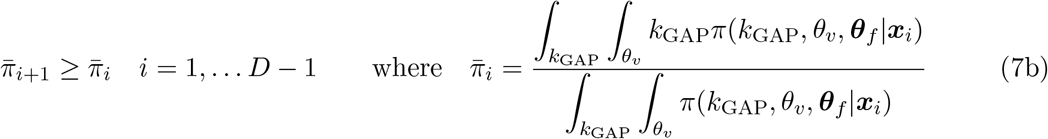

where *i* indexes the data sets (RhoGAP coexpression increases with *i*). Equation (7a) says that the maximum likelihood-to-evidence must be larger than 1 for at least one parameter set (*k*_GAP_, *θ*_*v*_), with ***θ***_*f*_ fixed, and (7b) says that the first moment of *k*_GAP_ must increase with RhoGAP coexpression (indexed by *i*). The posterior we show for a given (*k*_GAP_, *θ*_*v*_) is the total posterior *π*(*k*_GAP_, *θ*_*v*_, ***θ***_*f*_) summed over all ***θ***_*f*_ satisfying (7), divided by the number of such ***θ***_*f*_. This amounts to marginalizing only over fixed parameters that could reasonably fit all five data sets.

To select a particular fixed parameter set ***θ***_*f*_ for plotting (Fig. 8(C,D)), we identify fixed parameters that always increase the first moment 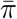 by a substantial amount (quantified by the *minimum* change in first moment), and have a high probability for all data sets (quantified by the *minimum* peak probability). The parameter set is therefore chosen by maximizing the score function

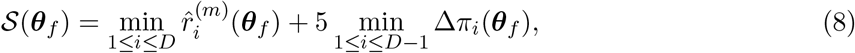

where the factor of 5 is chosen empirically to balance the two terms.

#### Bayes factor analysis

We use the Bayes factor, which measures the relative probabilities of two models, to compare the model where the filament lifetime *T*_fil_ varies with the GAP strength to the model where the basal nucleaton rate *q*_*b*_ varies with the GAP strength (Fig. 8). To compute a Bayes factor, we compute the total evidence for varying parameter *θ*_*v*_ by integrating over fixed parameter sets ***θ***_*f*_ which satisfy (7),

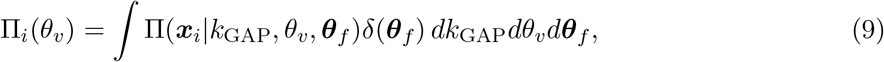

where *δ*(***θ***_*f*_) = 1 if (7) are satisfied and zero otherwise. Substituting *θ*_*v*_ = *T*_fil_, *q*_*b*_ into (9) and taking the ratio gives *B*_*i*_ = Π_*i*_(*q*_*b*_)*/*Π_*i*_(*T*_fil_), which is the Bayes factor comparing the model with varying *T*_fil_ to that with varying *q*_*b*_. A number greater than 1 favors the model in the numerator, and a number less than 1 favors the model in the denominator.

