## Supplemental tables and figures for "Learning activator–inhibitor dynamics at the cell cortex with neural likelihood ratio estimation"


| Parameter | Unit | Definition | Value/range |
|---|---|---|---|
| $k_{\mathrm{b}}$ | $1/(\mu\mathrm{m}^2\times\mathrm{s})$ | Basal Rho activation | 0.05 |
| $k_{\mathrm{ac}}$ | $1/(\mu\mathrm{m}^2\times\mathrm{s})$ | Autocatalysis | 1 |
| $K_{\mathrm{sat}}$ | $\mu\mathrm{m}^{-6}$ | Saturation | 0.1 |
| $D_\rho$ | $\mu\mathrm{m}^2/\mathrm{s}$ | Rho diffusivity | 0.1 |
| $k_{\mathrm{inac}}^{(0)}$ | 1/s | Basal Rho inactivation | 0.7 |
| $\nu_{p/d}$ | $\mu$m/s | F-actin growth/shrink rate | 1 |
| $g_w$ | $\mu$m | Filament thickness | 0.1 |
| $L$ | $\mu$m | Simulated domain size | 20 |
| $\Delta t$ | s | Time step size | 0.1 |
| $\Delta x$ | $\mu$m | Grid spacing | 0.2 |
| $k_{\mathrm{GAP}}$ | $\mu\mathrm{m}^2/\mathrm{s}$ | GAP strength | $[0.2, 0.6]$ |
| $T_{\mathrm{fil}}$ | s | Filament lifetime | $[1, 60]$ |
| $\ell_{\mathrm{max}}$ | $\mu$m | Max actin length | $[0.1, 10]$ |
| $f_b$ | $1/(\mu\mathrm{m})$ | Basal F-actin density | $[0, 8]^*$ |
| $f_\rho$ | $1/(\mu\mathrm{m})$ | Rho-mediated F-actin density | $[0, 200]$ |

**Table S1:** Model parameters. Top section lists fixed physical parameters, middle section lists fixed numerical parameters, and bottom section lists varied parameters. All of these are taken from previous work [30], with the exception of $k_{\mathrm{inac}}^{(0)}$ (reduced slightly to make it easier to sustain excitation) and $\nu_{p/d}$ (fixed in accordance with experimental measurements [62, 92]). As in prior work, the "thickness" of an F-actin filament is set to be much larger than the filament diameter ($g_w = 0.1$ $\mu$m), in order to resolve the filaments on a grid of reasonable size. Fig. S1 shows an example of the filament field $f(x)$ after it has been convolved with a Gaussian. *Samples are only collected on $f_b \in [0, 1.6/k_{\mathrm{GAP}}]$, as empirical testing revealed $f_b \geq 1.6/k_{\mathrm{GAP}}$ is too much F-actin to sustain excitation.

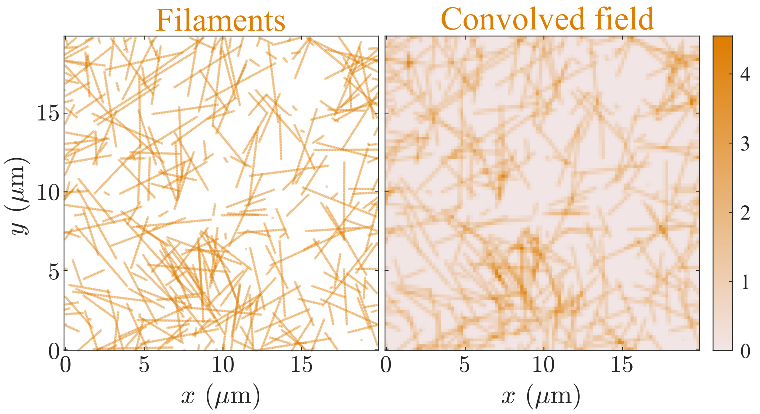


**Figure S1:** Sample of a filament field (left) and the result $f(x)$ when filaments are convolved with a Gaussian of width $g_w = 0.1$ $\mu$m (right).

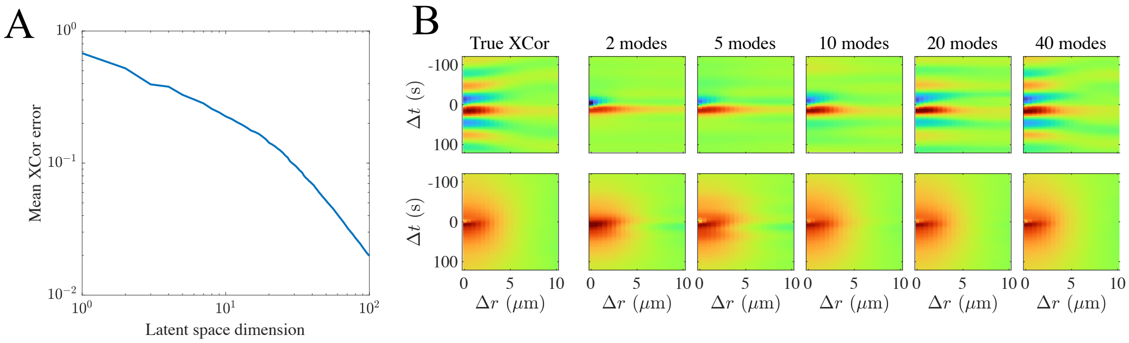


**Figure S2:** Compressing cross correlation by PCA. (A) The mean relative error in cross correlation (in the Frobenius norm) with varying numbers of PCA modes (using a training set of 10,000 cross-correlation functions and a testing set of 1000). (B) Example of compression on wave-like (top row) and pulse-like (bottom row) cross-correlation data. The minimal number of modes needed to capture the cross correlation functions is as low as 2 for a diffuse, pulse-like cross correlation, while about 20 modes are required for the wave-like cross correlation. Statistically, the mean error drops below 10% when 30 PCA modes are used, but 40 modes are required to reproduce the spatial structure of the wave-like cross correlation. We find that we can use many fewer modes for inference than are required to accurately resolve the cross correlation functions, in part because the strength of the first few modes often implies the rest.

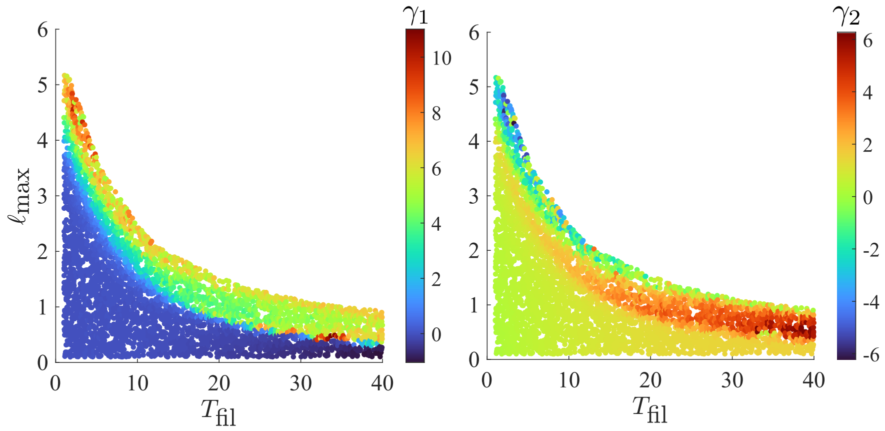


**Figure S3:** Samples of the model from two-dimensional ($T_{\text{fil}}, \ell_{\max}$) space. We set $k_{\text{GAP}} = 0.4$, $q_b = 0.054$, and $q_\rho = 0.86$, and scan over $T_{\text{fil}}$ and $\ell_{\max}$. Samples are colored by the first ($\gamma_1$, left) and second ($\gamma_2$, right) coordinate of the cross correlation in PCA space. While the coordinates $(\gamma_1, \gamma_2)$ are sufficient to infer an approximate region in parameter space, there is not an exact map between data and parameters, and there is more variation in the PCA modes in some regions of parameter space (e.g., top left) than others (e.g., bottom right). This implies (as shown more rigorously in Fig. S4) that the likelihood $p(\boldsymbol{x}|\boldsymbol{\theta})$ is broader in some regions of parameter space than others.

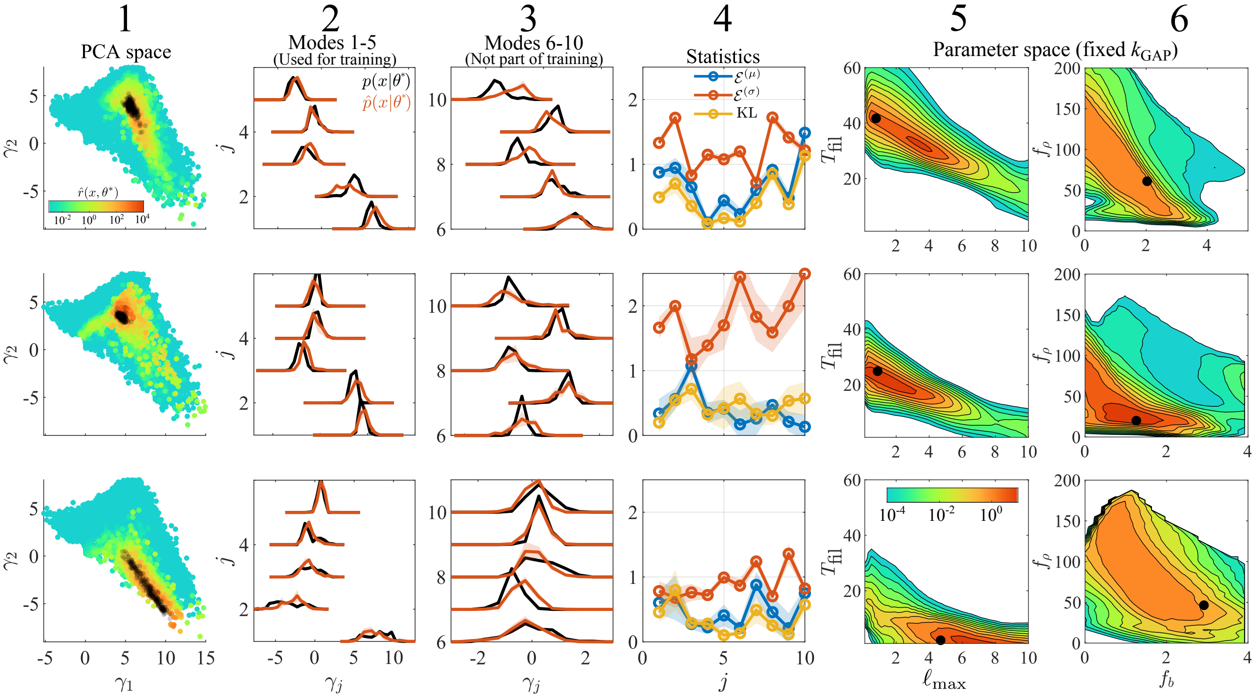


**Figure S4:** Inference on synthetic data is robust to different regions of parameter space. These are the same plots as Fig. 3, but for three different parameter sets (rows). Column 1: Samples from $p(\boldsymbol{x}|\boldsymbol{\theta}^*)$ in the first two components $(\gamma_1, \gamma_2)$ of PCA space (black), compared to likelihood ratio estimates $\hat{r}(\boldsymbol{x}, \boldsymbol{\theta}^*)$ for random samples from $p(\boldsymbol{x})$. Note how the spread of the true likelihood changes for the different parameters, and how this captured by the estimator. Column 2: One-dimensional histograms of distributions $p(\boldsymbol{x}|\boldsymbol{\theta}^*)$ (black) and $\hat{p}(\boldsymbol{x}|\boldsymbol{\theta}^*) = p(\boldsymbol{x})\hat{r}(\boldsymbol{x}, \boldsymbol{\theta}^*)$ in PCA space for the first five modes used to train the classifier. Column 3: Same histograms, but for modes 6–10 (not used to train the classifier). Column 4: Summary statistics (4) comparing the one-dimensional distributions for each mode. The relative error in the mean (blue), ratio of standard deviations (red), and KL divergence (yellow) are shown. Columns 5 and 6: Two-dimensional projections of the posterior $p(\boldsymbol{\theta}|\langle \boldsymbol{x}(\boldsymbol{\theta}^*)\rangle)$, where the black dots mark the true parameters $\boldsymbol{\theta}^*$ and $k_{\mathrm{GAP}}$ is fixed to the true value (from top to bottom, $k_{\mathrm{GAP}} = 0.3, 0.4, 0.4$) to make the F-actin densities $(f_b, f_\rho)$ meaningful. All results use 3 hidden layers with 20, 40, and 20 neurons each and $\lambda = 10^{-4}$ .

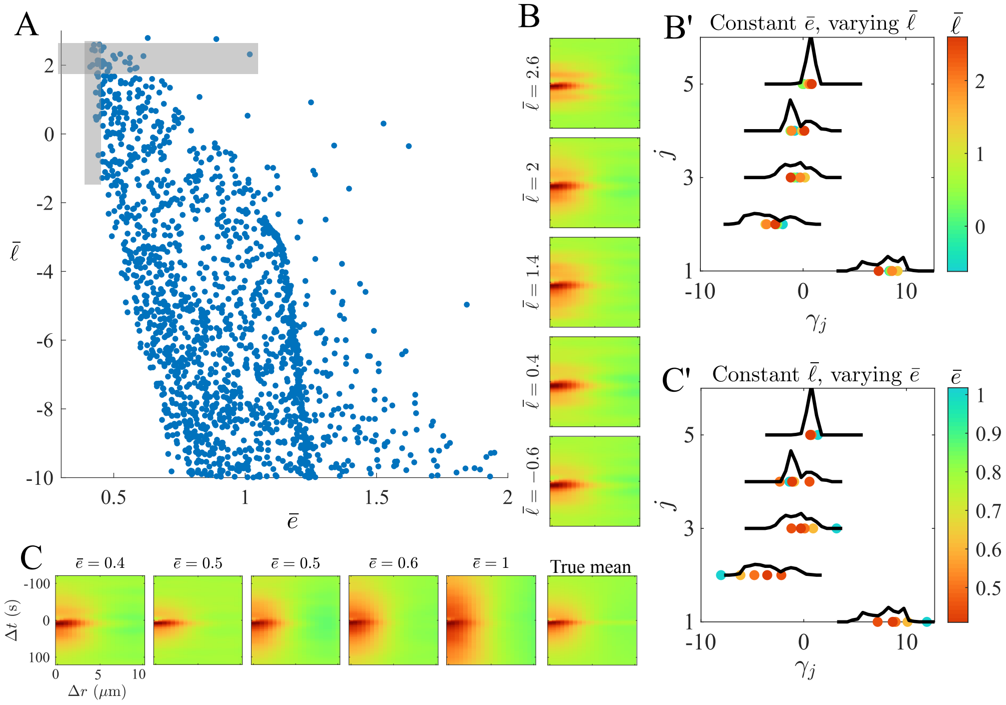


**Figure S5:** Likelihood ratio and cross correlation error contain complementary information. (A) Scatter plot comparing the mean error in cross correlation $\bar{e}(\boldsymbol{x}) = \left(\langle\|\boldsymbol{x} - \boldsymbol{x}(\boldsymbol{\theta})\|_F^2\rangle / \langle\|\boldsymbol{x}(\boldsymbol{\theta})\|^2\rangle\right)^{1/2}$ with the mean likelihood ratio $\bar{\ell} = \log_{10}\langle\hat{r}(\boldsymbol{x}, \boldsymbol{\theta})\rangle$. Gray rectangles denote regions (size 0.1 in cross correlation and 1 in log likelihood space) where there is variation in one quantity but not the other. (B) Samples of constant cross-correlation error and varying log likelihood. (B') Corresponding samples in PCA space, colored by the log likelihood, and compared to the true likelihood $p(\gamma_j|\boldsymbol{\theta}^*)$. (C) Samples of constant log likelihood but varying cross-correlation error. (C') Corresponding samples in PCA space, colored by the cross-correlation error, and compared to the true likelihood $p(\gamma_j|\boldsymbol{\theta}^*)$. The true mean cross correlation $\langle p(\boldsymbol{x}|\boldsymbol{\theta}^*)\rangle$ is shown at the bottom of (B) and right of (C). These are the same plots as Fig. 4, but for $\boldsymbol{\theta}^*$ we use the bottom parameter set in Fig. S4.

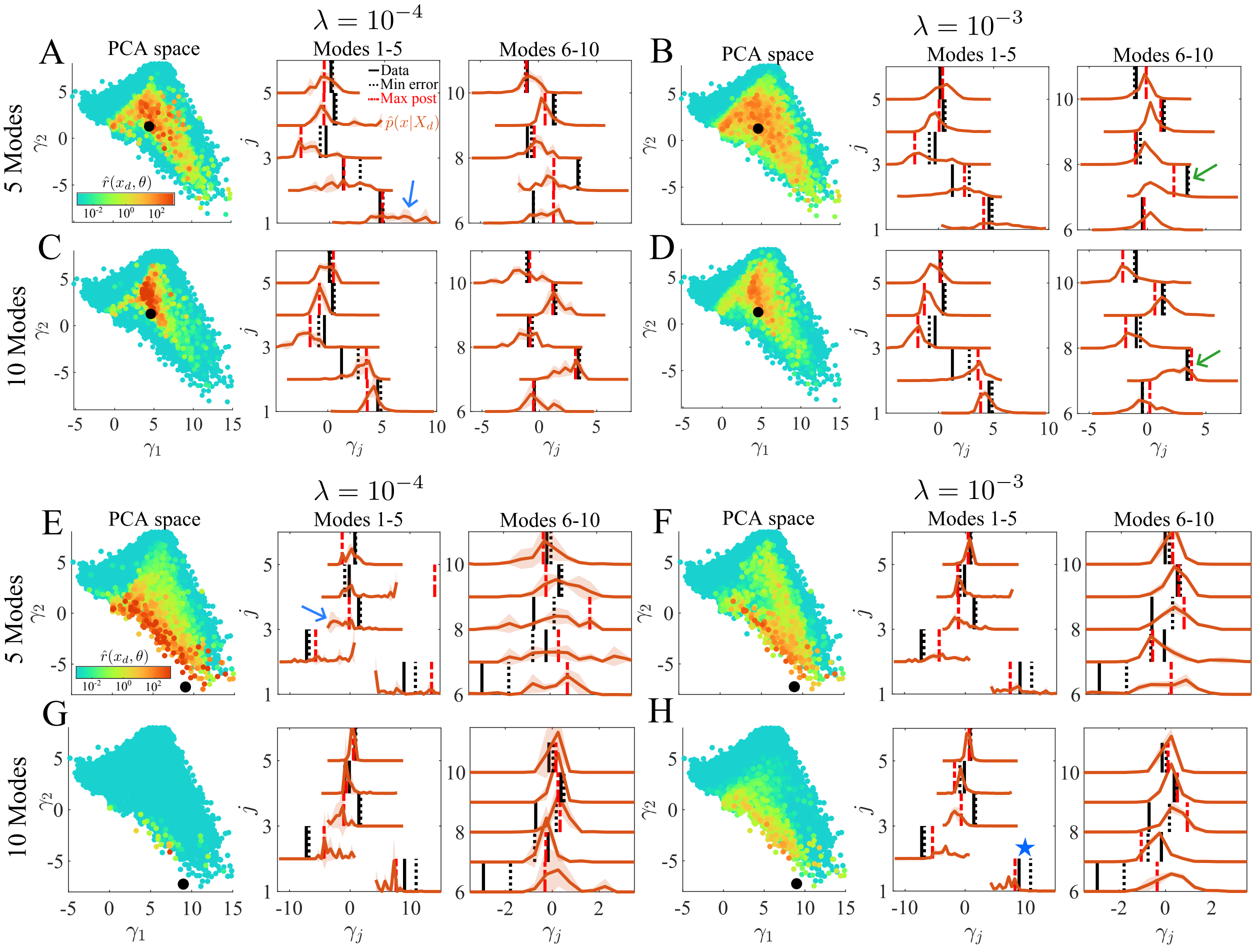


**Figure S6:** Pseudo-probability distributions $p(\boldsymbol{x}|\boldsymbol{X}_d)$ for the (A–D) starfish and (E–H) worm data with different classifier training settings. First column of each panel: Samples from $p(\boldsymbol{x}|\boldsymbol{\theta})$ in $(\gamma_1, \gamma_2)$ space, colored by $\hat{r}(\boldsymbol{X}_d, \boldsymbol{\theta})$ for 20,000 different values of $\boldsymbol{\theta}$ (from the training set). Experimental data are shown with a black dot. Second and third columns: Effective one-dimensional distributions $\hat{p}(\boldsymbol{x}|\boldsymbol{X}_d) = p(\boldsymbol{x}|\boldsymbol{\theta})\hat{r}(\boldsymbol{X}_d, \boldsymbol{\theta})$ in PCA space, compared to the true data (solid black lines), cross correlation with lowest error from the training set (dotted black lines), and average cross correlation from the maximum-posterior parameters (the dotted red line is an average over two samples). (A, E) Default classifier parameters ($\lambda = 10^{-4}$, 5 PCA modes) from synthetic validation. Blue arrows mark regions where the variance in the classifier predictions (estimated by computing $\hat{p}(\boldsymbol{x}|\boldsymbol{X}_d)$ a total of four times with two different classifiers) is notably high. (B, F) Additional regularization ($\lambda = 10^{-3}$) with 5 PCA modes. (C, G) More (10) PCA modes, with regularization held at $\lambda = 10^{-4}$. (D, H) More PCA modes and additional regularization ($\lambda = 10^{-3}$, 10 modes). Green arrows in (D) mark improvement of the classifier prediction relative to (B). Blue star in (H) marks where the maximum-posterior average is closer to the data than the minimum-error point.

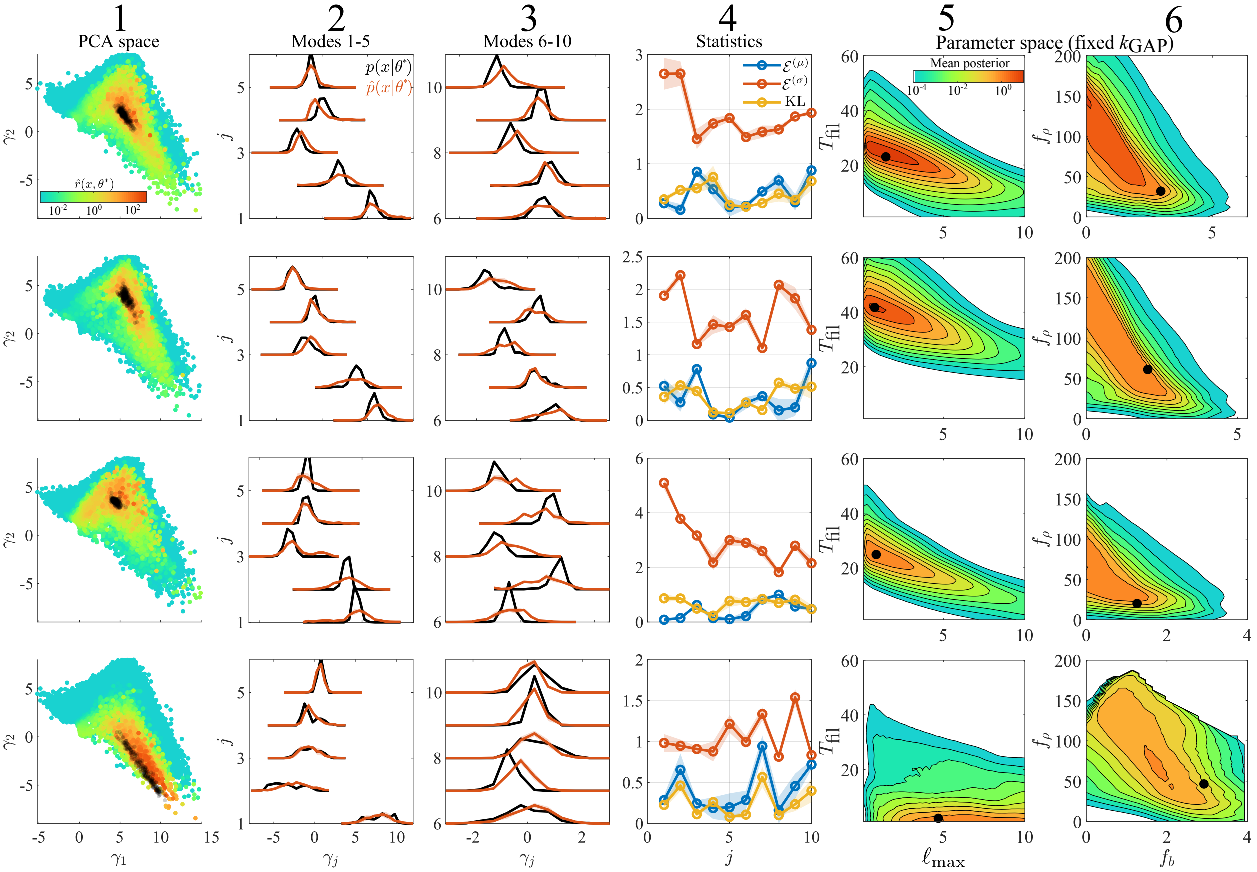


**Figure S7:** Inference on synthetic data using classifier settings for experimental data. These are the same plots as Fig. 3 and S4, but using a classifier with 3 hidden layers with 20, 40, and 20 neurons, $\lambda = 10^{-3}$, and 10 PCA modes (optimal settings for experimental data in Fig. 5D,H). Column 1: Samples from $p(\boldsymbol{x}|\boldsymbol{\theta}^*)$ in the first two components $(\gamma_1, \gamma_2)$ of PCA space (black), compared to likelihood ratio estimates $\hat{r}(\boldsymbol{x}, \boldsymbol{\theta}^*)$ for random samples from $p(\boldsymbol{x})$. Columns 2 and 3: One-dimensional histograms of distributions $p(\boldsymbol{x}|\boldsymbol{\theta}^*)$ (black) and $\hat{p}(\boldsymbol{x}|\boldsymbol{\theta}^*) = p(\boldsymbol{x})\hat{r}(\boldsymbol{x}, \boldsymbol{\theta}^*)$ for modes 1–5 and 6–10 (both used to train the classifier). Column 4: Summary statistics (4) comparing the one-dimensional distributions for each mode. The relative error in the mean (blue), ratio of standard deviations (red), and KL divergence (yellow) are shown. Columns 5 and 6: Two-dimensional projections of the posterior $p(\boldsymbol{\theta}|\langle \boldsymbol{x}(\boldsymbol{\theta}^*)\rangle)$, where the black dot is the true parameters $\boldsymbol{\theta}^*$ and $k_{\mathrm{GAP}}$ (from top to bottom, $k_{\mathrm{GAP}} = 0.25, 0.3, 0.4, 0.4$) is fixed to the true value to make the F-actin densities $(f_b, f_\rho)$ meaningful.

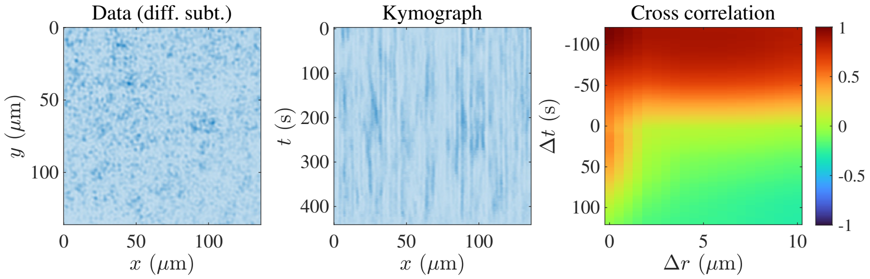


**Figure S8:** Corresponding data (Fig. 7) for no RGA coexpression (0 ng/$\mu$L). The trajectory is consistent with very low $k_{\mathrm{GAP}}$ levels, such that there are fluctuations around a single steady state of Rho. The cross-correlation function in this case resembles neither the model nor the other data sets.

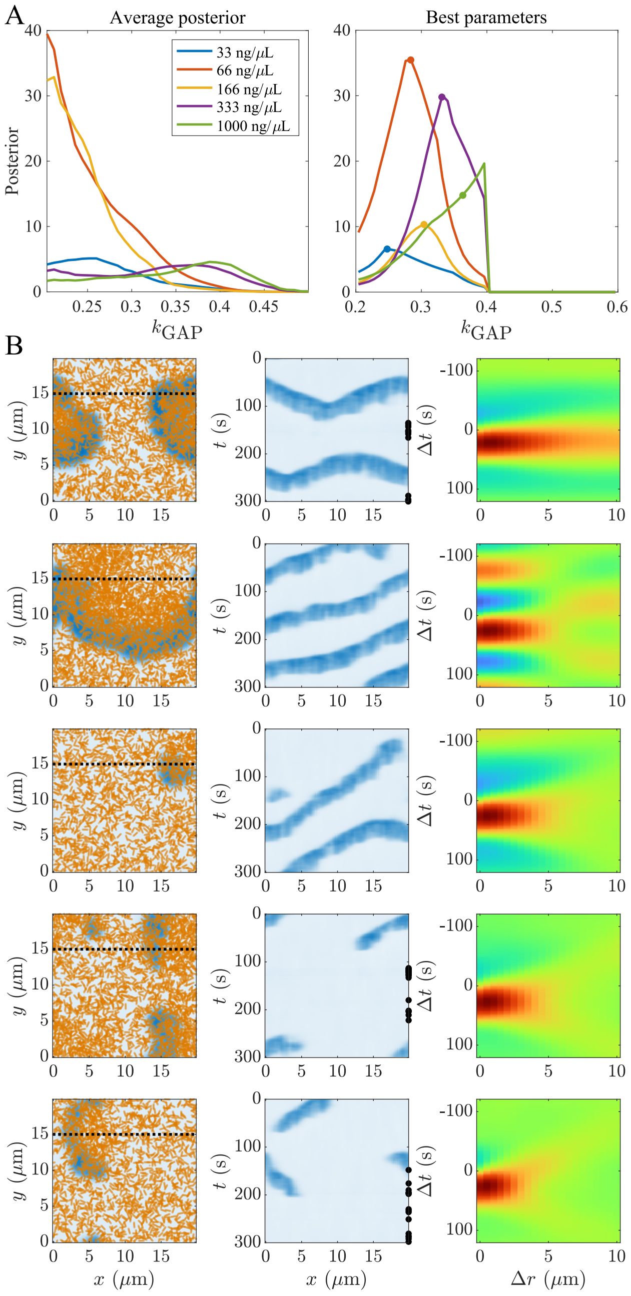


**Figure S9:** Best case inference when only $k_{\mathrm{GAP}}$ can vary. (A) Posterior density averaged over all parameters where the first moment of $k_{\mathrm{GAP}}$ increases with RGA expression (left) and over specific parameters $T_{\mathrm{fil}} = 51.6$, $\ell_{\max} = 0.706$, $f_b = 2.45$, $f_\rho = 24.5$ which optimize the balance between high probability and increasing first moment (right, this parameter set maximizes (8)). (B) Simulations from the dots in (A); from top to bottom, $k_{\mathrm{GAP}} = 0.252, 0.284, 0.308, 0.332, 0.36$. The last $k_{\mathrm{GAP}}$ value is adjusted to the maximum value that can sustain excitation for 120 s. Results are similar to Fig. 7.

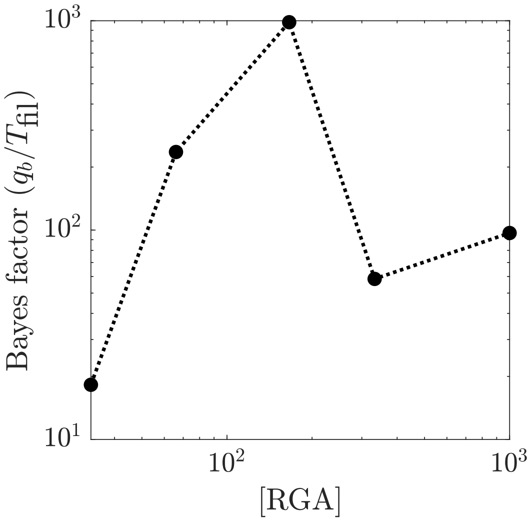


**Figure S10:** The Bayes factor comparing a model where only the basal nucleation rate $q_b$ varies with $k_{\rm GAP}$ to a model where only the filament lifetime $T_{\rm fil}$ varies with $k_{\rm GAP}$. For data point $i$, the Bayes factor is defined as $B_i = \Pi_i(q_b)/\Pi_i(T_{\rm fil})$, where the total evidence $\Pi_i$ is defined in (9).

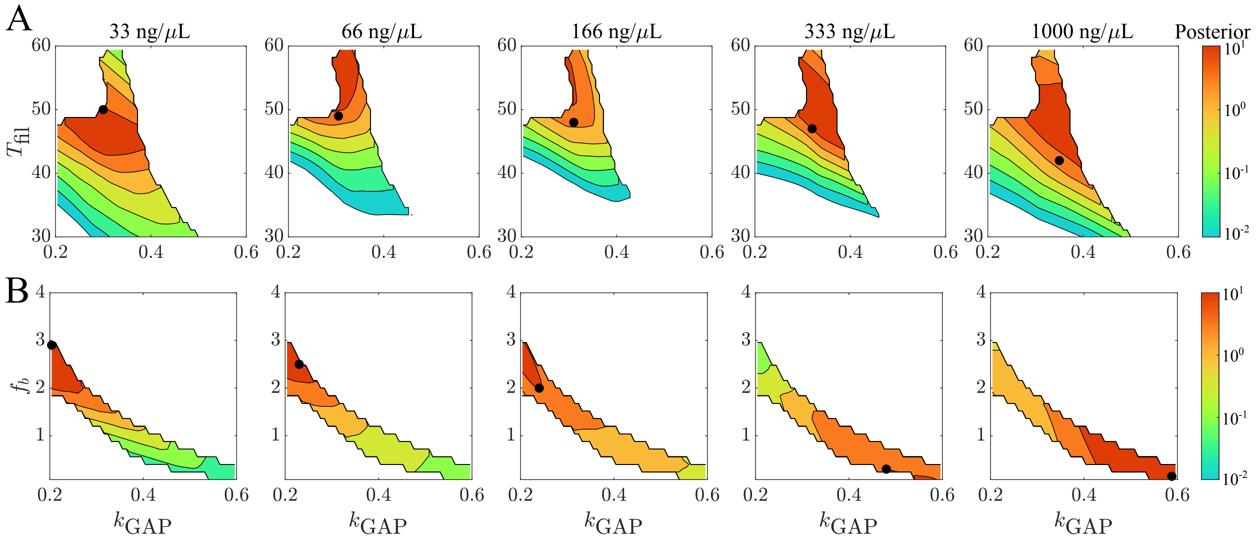


**Figure S11:** Posteriors for specific parameter sets that maximize (8). (A) Model where $T_{\rm fil}$ varies with $k_{\rm GAP}$. Black dots show the locations of samples chosen to decrease $T_{\rm fil}$ and increase $k_{\rm GAP}$ while remaining in high-likelihood regions (these are the same black dots as in Fig. 8A). (B) Model where $f_b$ varies with $k_{\rm GAP}$. Black dots show the locations of samples chosen to decrease $f_b$ and increase $k_{\rm GAP}$ while remaining in high-likelihood regions (these are the same black dots as in Fig. 8B). In this case only, it is possible to remain in regions with likelihood $10^1$.

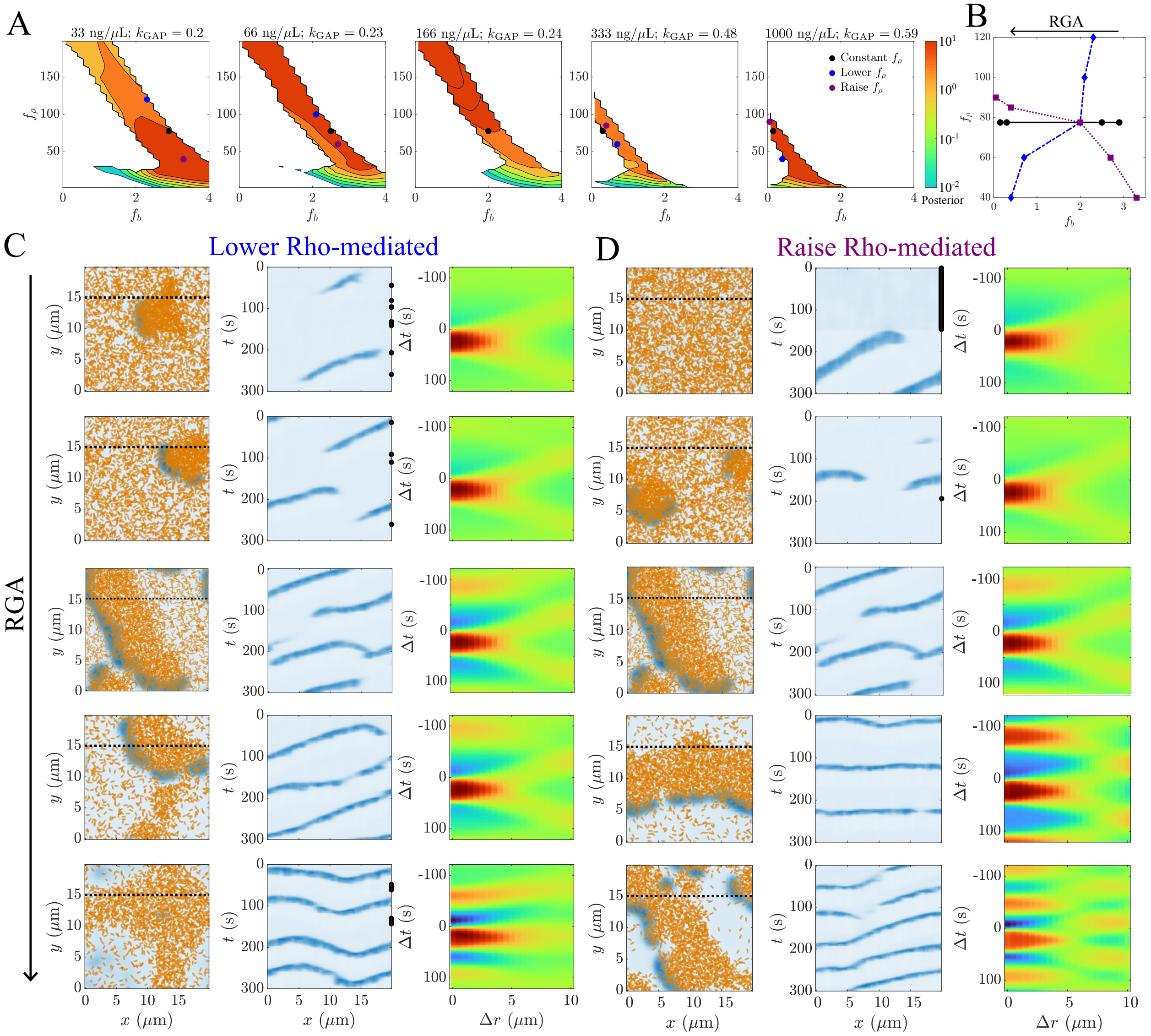


**Figure S12:** Rho-mediated nucleation can increase, decrease, or remain constant as GAP levels increase. (A) Posteriors for each RGA expression level in $(f_b, f_\rho)$ space with $k_{\text{GAP}}$ levels fixed as set in Fig. S11B. Black dots are samples when $f_\rho$ is constant, blue dots are samples when $f_\rho$ decreases with $k_{\text{GAP}}$, and purple dots are samples when $f_\rho$ increases with $k_{\text{GAP}}$. (B) Summary of parameter selections in $(f_b, f_\rho)$ space. RGA coexpression increases from right to left. (C) Samples from parameters with lower Rho mediated nucleation as $k_{\text{GAP}}$ increases (blue dots). (D) Samples from parameters with higher Rho mediated nucleation as $k_{\text{GAP}}$ increases (purple dots).
